# Developmental candidate GHP-88310/EIDD-3608 with high tolerability and oral efficacy in measles and respiratory paramyxovirus models

**DOI:** 10.64898/2025.12.21.695822

**Authors:** Carolin M Lieber, Josef D Wolf, Mugunthan Govindarajan, Jeong-Joong Yoon, Zachary M Sticher, Claire E Ruckel, Alexander I Leach, Lauren A Harrison, Dariia Vyshenska, Amalia A Cruz, Meghan K Andrews, Rebecca E Krueger, Robert M Cox, George R Painter, Alexander L Greninger, Michael G Natchus, Richard K Plemper

## Abstract

Orthoparamyxoviruses such as human parainfluenza virus type-3 (HPIV3) and measles virus (MeV), are a major health threat. We discovered an orally efficacious broad-spectrum inhibitor of orthoparamyxovirus polymerases. However, here we found that tolerability in higher mammals was limited. We report development of clinical candidate analog GHP-88310 (EIDD-3608), which combines improved oral efficacy with favorable tolerability in non-rodents (ferrets and dogs). GHP-88310 was active against HPIV3, Sendai virus (SeV), MeV, and related canine distemper virus (CDV). In 7-day tolerability studies, daily doses of 2,000 mg/kg were well tolerated. Pharmacokinetic analysis revealed altered plasma exposure of GHP-88310 compared to the original hit. In HPIV3-infected cotton rats, GHP-88310 lowered respiratory tract viral load. Dosing of ferrets infected with CDV, causing lethal measles-like disease, resulted in complete survival, reduction of viremia and shed viral load, and alleviated lymphocytopenia. Once-daily GHP-88310 was efficacious in the CDV-ferret and HPIV3-cotton rat models. The compound was sterilizing against HPIV3 at physiological concentrations in human airway epithelium organoids.

## Introduction

The orthoparamyxovirus subfamily contains major emerging and reemerging pathogens, such as measles virus (MeV), the human parainfluenzaviruses type 1 and 3 (HPIV1 and HPIV3), and the highly pathogenic Nipah and Hendra viruses (NiV and HeV) (*1, 2*). These viruses circulate in the human population or can readily transition species barriers as exemplified by spillover of NiV from *Pteropus* fruit bats to domestic animals and humans (*2*).

Although a live-attenuated measles vaccine has been available for decades, endemic transmission of MeV continues in large geographical areas including parts of Europe. Largely fueled by reports of an alleged link between the Measles, Mumps and Rubella (MMR) vaccine and the incidence of autism (*3, 4*), measles vaccination rates have declined in the last decade in the US. Further exacerbated by the COVID-19 pandemic (*5, 6*), low herd immunity in 2025 resulted in the largest measles outbreak in North America in 3 decades with over 1,500 confirmed cases around the epicenter in West Texas, three deaths, and thousands of additional cases in neighboring states, Mexico, and Canada (*7-9*). Approximately 20% of patients were hospitalized with severe disease (*10*). However, no approved antivirals for improved measles management beyond supportive care exist (*11-13*). A major obstacle that has hampered drug development against MeV is the challenge of conducting clinical efficacy trials in a predominantly pediatric patient population (*14-16*). In contrast, related HPIV3 can reinfect people throughout life. Older adults and the immunocompromised are at greatest risk of HPIV3 spread to the small airways, causing lower respiratory tract infection (LRTI) that presents with bronchiolitis and pneumonia (*17, 18*). Adult hematopoietic stem cell transplant recipients are among the most vulnerable groups with mortality rates from HPIV3 LRI of up to 75% (*19, 20*), and no vaccine prophylaxis or therapeutic is currently available to manage the disease.

We have identified HPIV3 as the primary indication for an anti-orthoparamyxovirus drug development program based on the rationale that the HPIVs constitute an unmet medical need with an estimated 3 million medically attended cases in the United States annually (*21, 22*); that progression to severe HPIV3 disease in at-risk adults is relatively slow with a median of 3 days between upper respiratory symptoms and onset of pneumonia, opening a window for intervention (*20*); and that precedence for clinical evaluation of HPIV3 inhibitors in transplant patients has been established (*19, 20*).

In previous work, we identified a novel non-nucleoside inhibitor class of the paramyxovirus RNA-dependent RNA polymerase (RdRP) complex through an anti-HPIV3 high-throughput drug screen (*23, 24*). The hit compound, GHP-88309, showed unusually broad-spectrum sub-micromolar activity against orthoparamyxoviruses of the respirovirus (HPIV1, HPIV3, SeV), morbillivirus (MeV and CDV), and henipavirus (Ghana henipavirus and Cedar virus) genera (*23-25*). Mechanistic characterization revealed that GHP-88309 binding prevents initiation of the viral RdRP complex at the promoter, most likely by sterically blocking structural reorganization of the polymerase from initiation into elongation configuration (*23, 24, 26*). For proof-of-concept, we established oral efficacy in the SeV-mouse model as a lethal substitute assay for human respirovirus disease (*23, 24*). In parallel, we validated broad anti-orthoparamyxovirus efficacy in CDV-infected ferrets, which develop measles-like disease that presents with high fever, rash, and lymphocytopenia (*25*). The infection recapitulates severe measles cases, since CDV pathogenicity in ferrets is high and all infected animals succumb within 10-12 days of infection (*25*). A course of oral GHP-88309 started after the onset of clinical signs but before peak viremia changed outcome and all treated animals survived (*23, 24*). Thus, GHP-88309 met three fundamental requirements of the target product profile that we require of an anti-orthoparamyxovirus candidate: orally bioavailable to facilitate rapid self-administration; active against different high-consequence orthoparamyxoviruses including an indication with a clear path to clinical testing; and shelf-stable under ambient conditions for stockpiling and rapid distribution. However, the envisioned clinical use in pediatric patients and the immunocompromised demands high tolerability with wide safety margins.

In this study, we report an unexpected tolerability limit of GHP-88309 in higher mammals of 50 mg/kg. In search of a clinical candidate, we launched a synthetic lead development program that yielded a novel chemotype class with favorable tolerability in rodent and non-rodent (ferrets and dogs) toxicology species. We report enhanced oral efficacy of this candidate against the primary HPIV3 indication in a relevant animal model, show a broadened therapeutic window against the secondary measles indication, and establish dosing paradigms against HPIV3 in disease-relevant human airway epithelium organoid cultures based on PK-informed physiologically achievable dynamic concentrations.

## Results

Prolonged 17-day treatment of CDV disease in ferrets with the first-generation lead GHP-88309 was highly effective at a minimally efficacious oral dose of 50 mg/kg twice daily (*bis in die* (b.i.d.)), and all animals survived the infection (*25*). In mice, multi-dose administration of GHP-88309 at 150 mg/kg (highest dose tested) b.i.d. was well tolerated without adverse events (*23*).

### Synthetic lead development to overcome tolerability limitations of GHP-88309

To estimate the therapeutic index (TI) of GHP-88309 in a non-rodent species, we increased oral dose levels to 150 mg/kg in a multi-dose PK study in search of tolerability limits in ferrets (Fig. 1A). Unexpectedly, ferrets in the 150 mg/kg group reached endpoint within hours following the second dose (Fig. 1B, Fig. S1A,B). Complete blood count (CBC) analysis revealed a drastic drop in platelet counts in animals of this group after compound administration (Fig. S1C). Plasma PK analysis of the 50 mg/kg group on day 7 (after 14 doses) showed high plasma concentration peaks (C_max_ 43.5 μM) and an overall exposure of 270 μM × hr (Fig. 1C). A single ascending dose study confirmed that a dose level of 50 mg/kg was well tolerated, whereas animals receiving 150 mg/kg showed transient paralysis and a drop in body temperature, and ferrets dosed at 500 mg/kg died within 4 hours of dosing (Fig. 1D-F).

**Figure 1.**
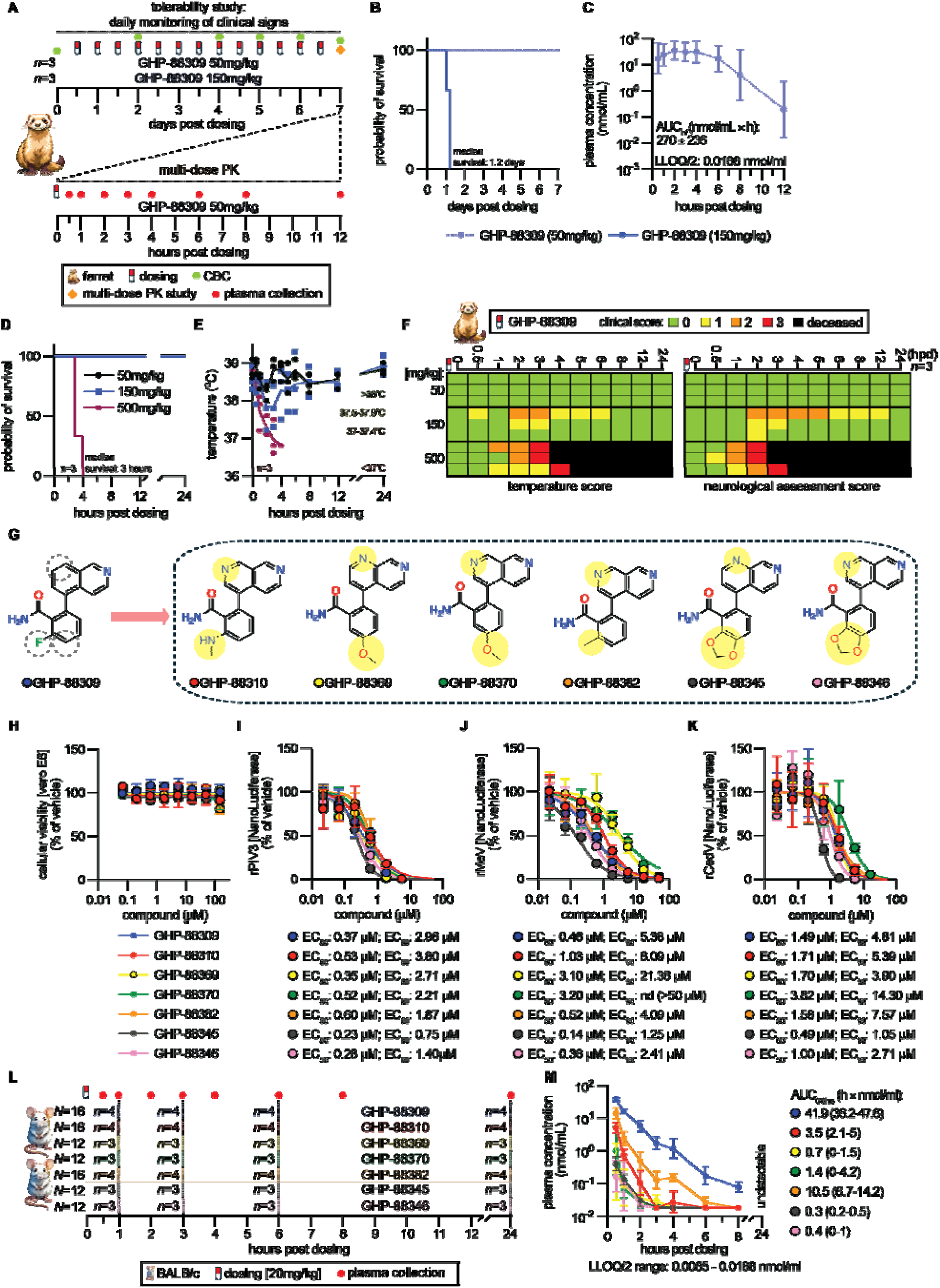
SAR development of broad-spectrum GHP-class polymerase inhibitors. **A)** Study schematic of multi-dose GHP-88309 tolerability and PK assessment. Ferrets received 14 oral doses of GHP-88309 at two different concentrations, administered twice daily. **B)** Survival curves of animals from (A). **C)** PK profile of GHP-88309 (50 mg/kg) in ferret plasma following 14 oral doses administered b.i.d. Exposure is specified. **D-F)** Survival curves (D), body temperature (E), and clinical scores (F) of animals after single oral dose of GHP-88309. Body temperature scoring: 0, >38°C; 1, 37.5-37.9°C; 2, 37-37.4°C; 3, <37°C. Neurologic scoring: 0, explores objects, weasel war dance, intact reflexes; 1, decreased interest in exploring objects; 2, lethargic, and/or altered gait, and/or decreased reflexes; 3, obtunded, and/or seizures, and/or paralysis in one or more limbs. **G)** Top-ranking GHP-88309 analogs. Dashed circles mark regions that QSAR models (*23*) have identified as most amenable to synthetic elaboration. **H)** Evaluation of cellular viability of GHP compounds on Vero-E6 cells. **I-K)** Recombinant virus reporter-based dose response inhibition of HPIV3 (I), MeV (J), and CedV (K), n ≥3. **L)** Study schematic of single-dose PK assessment in BALB/c mice. Mice received an oral dose of GHP-class (20 mg/kg) and were monitored over 24 hours. Blood was collected from each sub-group of mice twice, following euthanasia. **M)** Single-dose PK profile in mouse plasma. Exposure is given as AUC_0-8hrs_ with 95% confidence intervals in parenthesis. Group sizes (n) specified in study schematics (A,L). Log-rank (Mantel-Cox) test with median survival shown (B,D). Symbols represent geometric mean ± geometric standard deviation (SD) (C,M). Variable slope 4-parameter non-linear regression modeling (H-K), n ≥3. Symbols represent means ± SD. LLOQ/2, 50% of lowest limit of quantitation; AUC, area under the curve.

Building on our initial understanding of the structure-activity relationship (SAR) of the scaffold (*23*), we launched a synthetic lead development program with the primary objective to improve tolerability in non-rodent species. Six azaquinoline lead candidates emerged from this exercise (Table S1), each featuring a substituted heterocyclic ring (Fig. 1G). None showed appreciable cytotoxicity in cell culture (Fig. 1H) and all maintained antiviral potency *in vitro* within a <2-fold range against the HPIV3 primary indication (Fig. 1I) and respective <8-fold and <3-fold range against the morbillivirus (Fig. 1J) and henipavirus (Fig. 1K) secondary indications.

Single-dose PK prescreening of these six candidates in mice at an oral dose level of 20 mg/kg revealed highest plasma exposure for, in descending order, GHP-88382, GHP-88310, and GHP-88370. However, none of these analogs reached the high exposure levels of GHP-88309 (Fig. 1L). Based on PK performance, we selected these three analogs for proof-of-concept efficacy testing against the primary respirovirus indication.

### GHP-88310 efficacy against the primary respirovirus indication

Using the lethal SeV 129X1/SvJ mouse model as a surrogate for human respirovirus disease (*27*), we screened the three developmental candidates in comparison with GHP-88309, each dosed orally at 150 mg/kg b.i.d., initiated 24 hours after infection (Fig. 2A). All animals receiving GHP-88309 and GHP-88310 survived without developing clinical signs, whereas GHP-88382 mediated only partial survival and GHP-88370 provided no therapeutic benefit (Fig. 2B, Fig. S2A,B). Viral load in the upper (Fig. 2C) and lower (Fig. 2D) respiratory tract of GHP-88310-treated animals was significantly reduced compared to vehicle-treated mice. Effect size was equivalent to that achieved by GHP-88309. In contrast, GHP-88382 reduced viral burden only in the lower respiratory tract and GHP-88370 had no significant effect on viral load.

**Figure 2.**
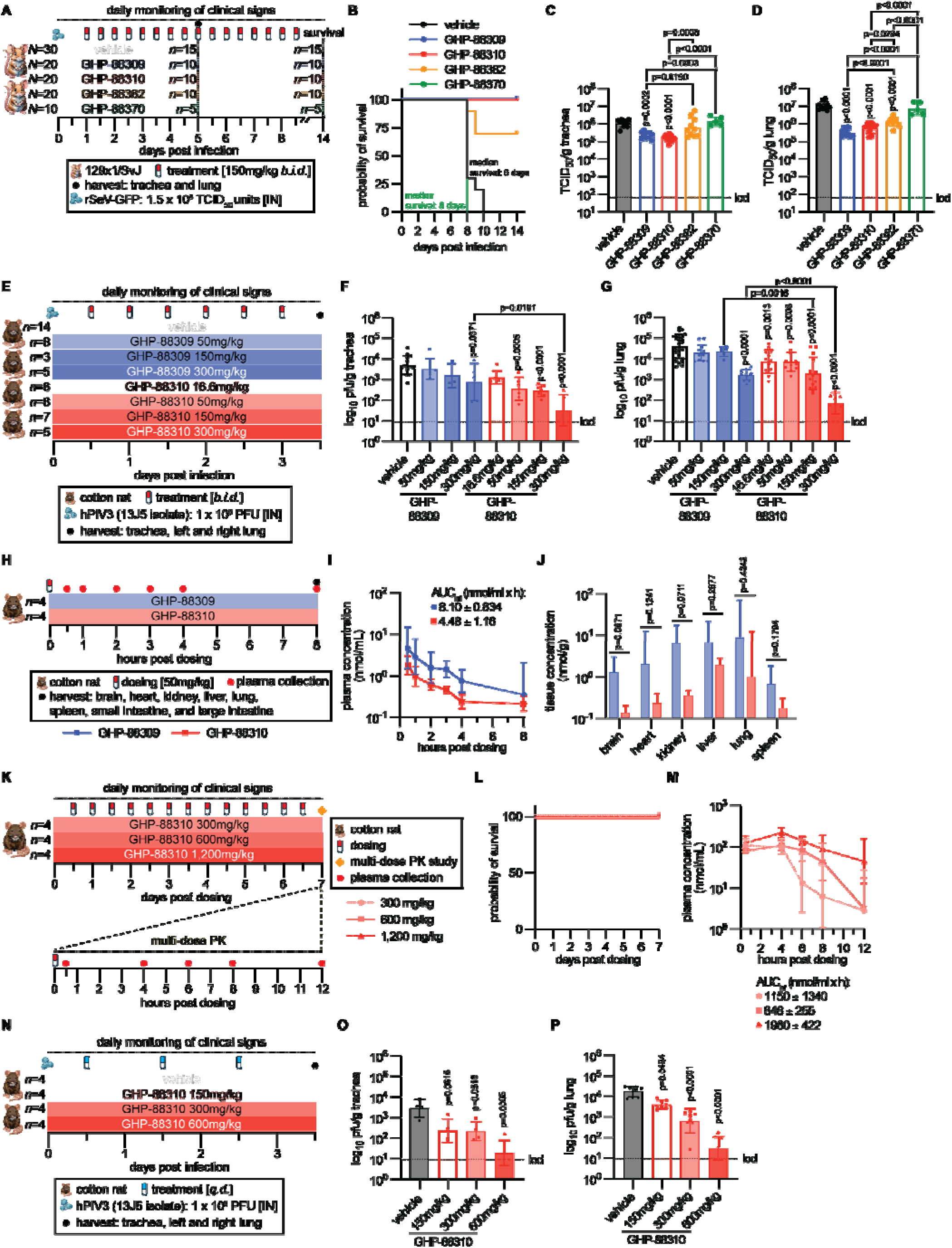
PK and efficacy evaluation of GHP-88309 analogs in rodents. **A)** Study schematic. 129x1/SvJ mice were infected intranasally with 1.5×10^5^ TCID_50_ units of recSeV. Compounds were administered orally b.i.d., starting 1 dpi and continuing through 8.5 dpi. **B-D)** Survival curves (B), trachea (C) and lung (D) viral titers determined 5 dpi. **E)** Study overview. Direct comparison of GHP-88309 and GHP-88310 efficacy conducted in cotton rats. Animals were intranasally infected with 1×10^6^ TCID_50_ units of HPIV3 isolate 13J5. Oral dosing was initiated 12 hours post infection (hpi) and continued b.i.d. **F,G)** Trachea (F) and combined left and right cranial lungs (G) were collected 3.5 dpi and viral titers determined by plaque assay. **H)** Study outline of single-dose PK and tissue distribution analysis in cotton rats, comparing 50 mg/kg doses of GHP-88309 and GHP-88310. **I,J)** PK profiles of GHP-88309 and GHP-88310 in cotton rat plasma (I) and tissue concentrations of GHP-88309 and GHP-88310 measured 8 hours post dosing (J). Unpaired, two-tailed t-test. Bars indicate the geometric mean ± geometric SD. **K)** Multi-dose GHP-88310 tolerability and PK study. Cotton rats received 14 oral doses, administered b.i.d. **L,M)** Survival curves (L) and PK profile (M) after 14 doses of GHP-88310. **N)** Overview of efficacy evaluation of GHP-88310 administered q.d. at different concentrations in cotton rats. Animals were intranasally infected with 1×10^6^ TCID_50_ units of HPIV3, and oral dosing started 12 hpi. **O,P)** Trachea (O) and combined left and right cranial lung lobes (P) were collected 3.5 dpi and viral titers determined by plaque assay. In (B,L), log-rank (Mantel-Cox) test with median survival. In (C,D,F,G,O,P), bars indicate the geometric mean ± geometric SD, symbols represent individual biological repeats. One-way ANOVA with Tukey’s (between group comparisons) or Dunnet’s (comparisons to vehicle) post-hoc multiple comparisons test. In (I,M), symbols represent geometric means ± geometric SD.

Accordingly, we selected GHP-88310 for a dose-to-failure study against the primary indication in the HPIV3 cotton rat model, measuring viral load in the respiratory tract 3.5 days post-infection (dpi; Fig. 2E). In this model, the minimal dose of original hit GHP-88309 that significantly reduced viral load in both trachea and lungs was 300 mg/kg b.i.d. (Fig. 2F,G; Fig. S3A,B). GHP-88310 was substantially more potent, showing lowest efficacious doses of 50 mg/kg b.i.d. and 16.5 mg/kg b.i.d. in the upper and lower respiratory tract, respectively. Plasma exposure of GHP-88310 in cotton rats was approximately one-half that of GHP-88309 after a single oral dose of 50 mg/kg, despite greater efficacy of GHP-88310 (Fig. 2H,I). Corresponding variations in tissue distribution were greatest in brain, heart, and kidneys, smallest in liver and lung (Fig. 2J). However, none of these differences were statistically significant.

A 7-day ascending multi-dose study with GHP-88310 culminating in a 2,400 mg/kg daily dose (Fig. 2K) showed that the compound was well tolerated at all dose levels without changes in bodyweight, temperature, or other adverse signs (Fig. 2L; Fig. S4A,B). Plasma exposure built up to nearly 2,000 μM × hr at the highest dose and plasma exposure levels reached by animals in the highest (1,200 mg/kg) and second highest (600 mg/kg) dose groups were near dose-proportional (1,980 vs 848 nmol/ml × h). (Fig. 2M). Trough plasma concentrations at the lowest dose tested exceeded 19 μM, corresponding to 5-times the cell-culture EC_90_ concentration of GHP-88310 against HPIV3. Based on these exposure levels, we examined anti-HPIV3 efficacy in a once-daily (*quaque die* (q.d.)) therapeutic dosing regimen, starting treatment of cotton rats 12 hpi (Fig. 2N). Oral GHP-88310 at 150 mg/kg q.d. or higher significantly reduced viral load in trachea and the lower respiratory tract (Fig. 2O,P) without changes in body weight or temperature (Fig. S5A,B), which was consistent with a direct correlation between plasma exposure and *in vitro* EC_90_ concentrations.

### GHP-88310 tolerability in non-rodent toxicology species

Having validated oral efficacy of GHP-88310 against the primary indication in cotton rats, we explored PK properties of the compound in ferrets in a single ascending dose oral PK study (Fig 3A). A dose level of 500 mg/kg GHP-88310 was well tolerated and plasma exposure increased dose-proportionally between 150 and 500 mg/kg doses (Fig. 3B). A subsequent 7-day ascending multi-dose tolerability study in ferrets demonstrated that administering GHP-88310 at a daily dose of up to 2,000 mg/ml (Fig. 3C) had no negative effect on bodyweight, body temperature, and CBC parameters including platelet counts (Fig. S6A-E). All animals survived the full 7-day course without signs of adverse effects, in contrast to reference animals dosed with GHP-88309 at 150 mg/kg, which succumbed within hours of receiving the second dose (Fig. 3D). Plasma concentrations examined once-daily 4 hours after dosing (expected C_max_) plateaued dose-dependently after administration of four (500 mg/kg) to eight (1,000 mg/kg) doses (Fig. 3E). Overall exposure after 7 days (14 doses) increased dose-dependently from 336 μM × hr (50 mg/kg b.i.d.) to 2,140 μM × hr (500 mg/kg b.i.d) (Fig. 3F). At a dose of 50 mg/kg b.i.d., total plasma exposure of GHP-88310 was slightly higher than that of GHP-88309 with 336 μM × hours versus 270 μM × hours at study end (Fig. 1C vs Fig. 3F). Furthermore, tissue distribution, determined 12 hours after the last of 14 doses administered, showed good dose-dependency. Average GHP-88310 tissue concentration was approximately 10 nmol/g (150mg/kg group) in all soft organs analyzed including ferret brain (Fig S7).

**Figure 3.**
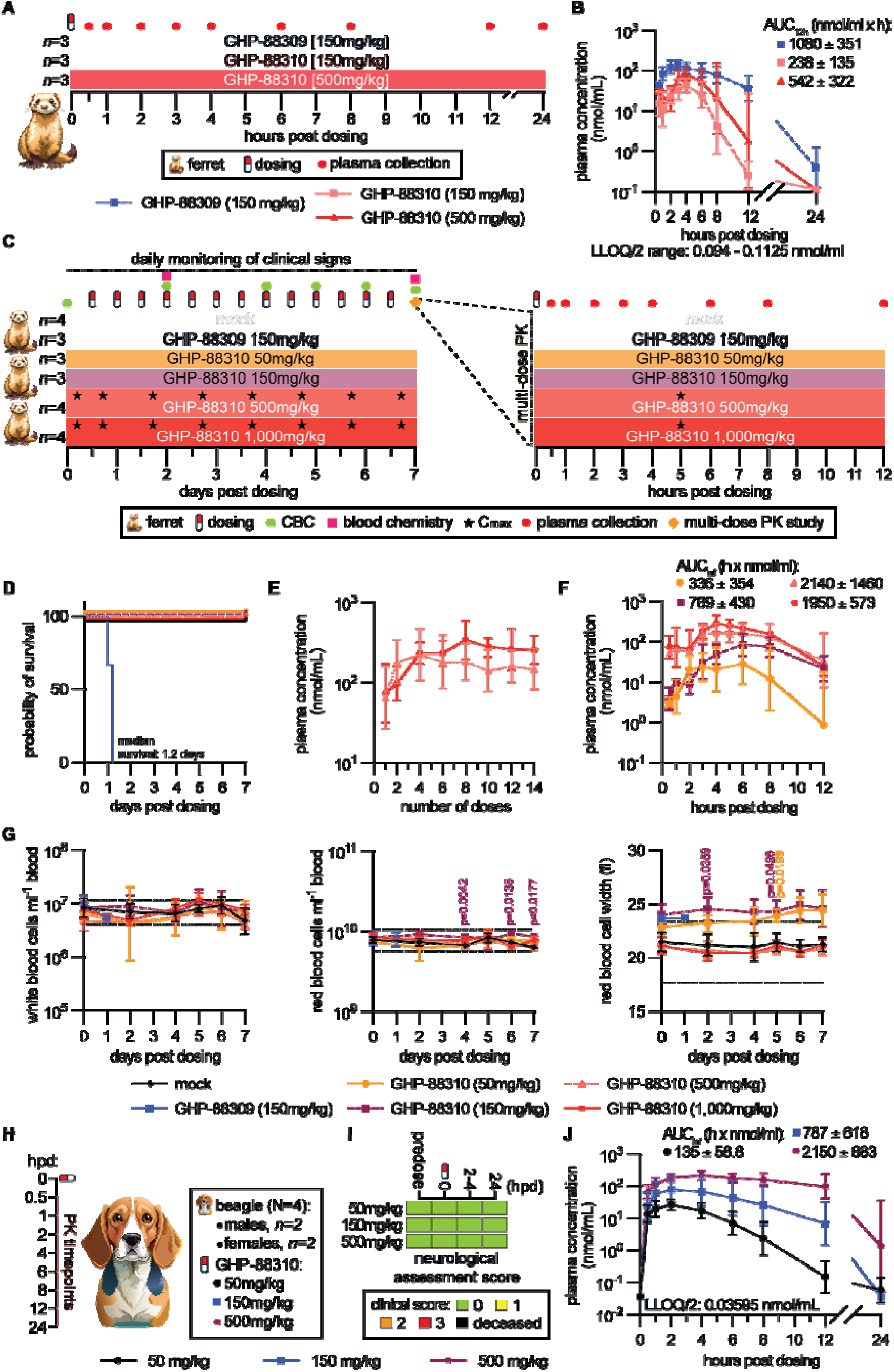
Single and multi-dose tolerability and PK of twice daily GHP-88310 in ferrets. **A)** Overview of single oral dose PK study of GHP-88309 and GHP-88310 in ferrets. **B)** PK profile of GHP-88309 and GHP-88310 in ferret plasma. **C)** Schematic of multi-dose tolerability and PK study of GHP-88309 and GHP-88310 at different concentrations. Ferrets received 14 oral doses, administered b.i.d. Maximal plasma concentration (C_max_) was measured daily in ferrets administered 500 and 1,000 mg/kg GHP-88310. **D)** Survival curves; data for the 150 mg/kg GHP-88309 group, originally shown in (Fig. 1A-B, are included for direct comparison. Log-rank (Mantel-Cox) test with median survival. **E)** C_max_ of GHP-88310 to monitor plasma build-up after doses 1, 2, 4, 6, 8, 10, 12, and 14. **F)** PK profile of GHP-88310 in ferret plasma after 14 oral doses. **G)** CBC analysis of ferret blood samples quantifying white blood cells (left), red blood cells (middle), and red blood cell width (right). Normal range for each parameter is shown in green shading. (Left, middle), symbols indicate the geometric mean ± geometric SD, (right) symbols represent means ± SD. Two-way ANOVA followed by Dunnet’s (comparisons to mock) post-hoc multiple comparisons test. **H-J)** Single dose PK study of GHP-88310 in dogs. Shown are study outline (H), neurologic status scores (I) (0, explores objects, intact reflexes; 1, decreased interest in exploring objects; 2, lethargic, and/or altered gait, and/or decreased reflexes; 3, obtunded, and/or seizures, and/or paralysis in one or more limbs), and PK profile of GHP-88310 in dog plasma (J). In (B, E-F, J), symbols indicate geometric means ± geometric SD. LLOQ/2, 50% of lowest limit of quantitation.

CBC parameters of GHP-88310-treated animals did not deviate from the normal range over the course of the study (Fig. 3G) and whole blood chemistry panels taken on days 2 and 7 were likewise unremarkable (Fig. S8). In contrast, a reference animal that had received two doses of GHP-88309 at 150 mg/kg showed changes in three of four liver enzymes examined – alkaline phosphatase (ALP), alanine transaminase (ALT), and aspartate aminotransferase (AST) – by more than 750% one day post-dosing and prior to death, suggesting GHP-88309-induced hepatotoxicity. Due to rapid death after repeated administration of GHP-88309 at this dose level, a comprehensive blood chemistry analysis of all animals in the reference group was not possible.

Based on greatly improved tolerability of GHP-88310 compared to GHP-88309, we advanced the compound to single ascending-dose PK testing in beagle dogs, the predominant non-rodent toxicology species used to inform investigational new drug (IND)-enabling safety studies (Fig. 3H). Dogs were dosed orally with GHP-88310 at 50, 150, or 500 mg/kg, each followed by assessment of plasma exposure over a 24-hour period (Fig. 3H). All dose levels were well tolerated without adverse signs or weight loss of the animals (Fig. 3I). Plasma exposure ranged from 135 to 2,150 μM × hr (Fig. 3J), demonstrating approximate dose-proportionality and exceeding at high-dose that observed in ferrets. These results demonstrated that GHP-88310 plasma exposure levels exceeding those at which GHP-88309 was lethal in ferrets were well tolerated in both ferrets and dogs (Fig. 3B and 3J).

### GHP-88310 efficacy against measles-like disease in ferrets

Measles has emerged as an urgent secondary indication for GHP class paramyxovirus inhibitors due to growing vaccination hesitancy in many high-income countries and endemic MeV transmission in large geographical regions in sub-Saharan Africa and Southeast Asia (*12, 28, 29*). We have demonstrated potent oral efficacy of GHP-88309 against the CDV/ferret surrogate model of human measles (*25*). GHP-88310 and GHP-88309 had virtually identical antiviral activity against both MeV (Fig. 1J) and CDV (Fig. 4A, Fig. S9) in cell culture, returning active (EC_50_) concentrations within a 2-fold range of each other.

**Figure 4.**
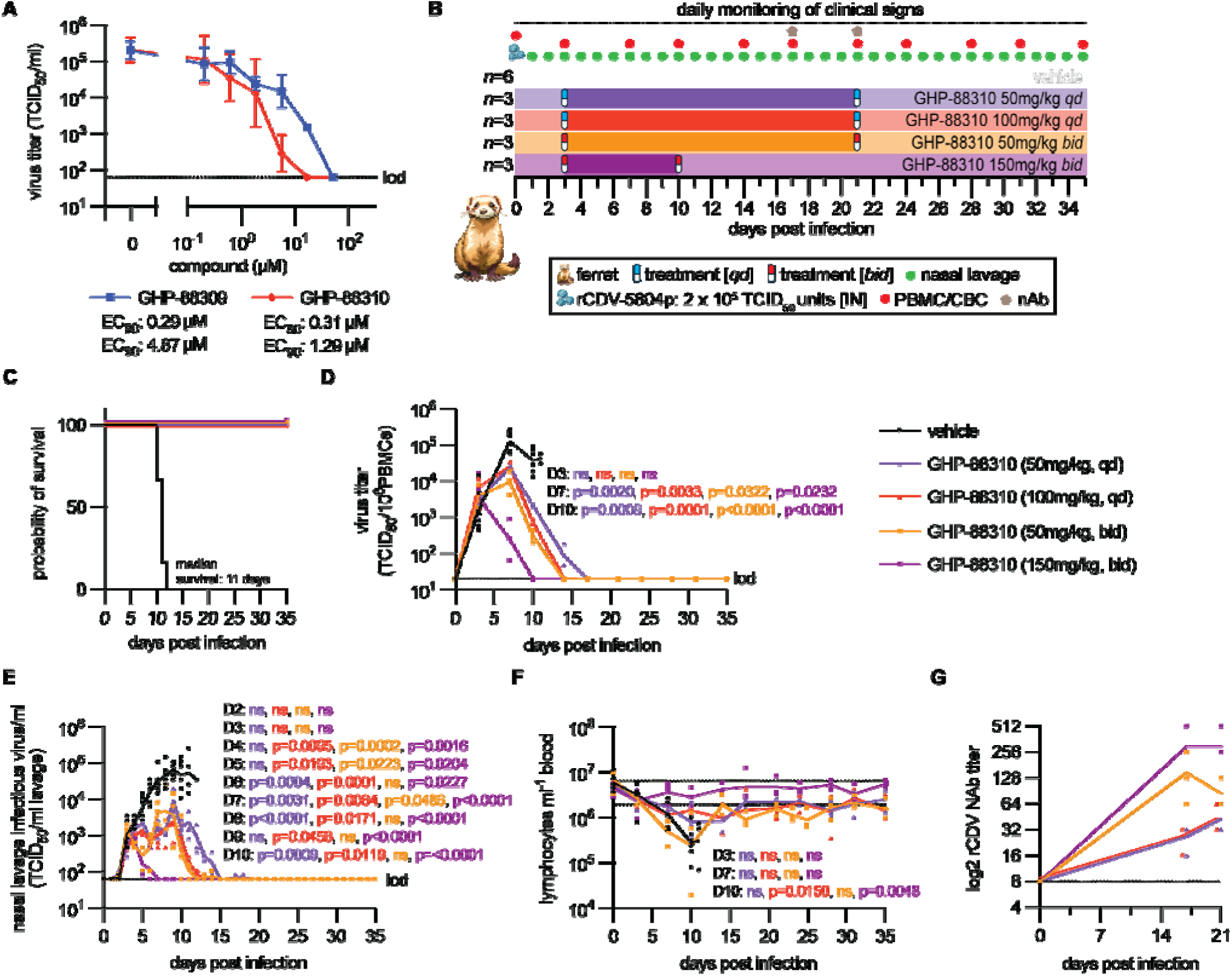
Therapeutic effect of GHP-88310 on CDV infection in ferrets. **A)** Virus yield reduction of recCDV-5804p. Variable slope 4-parameter non-linear regression modeling; n ≥3. Symbols indicate the geometric mean ± geometric SD. **B)** Overview of assessment of q.d. and b.i.d. GHP-88310 therapeutic potential. Ferrets were infected intranasally with 2×10^5^ TCID_50_ units of recCDV-5804p. Oral administration of GHP-88310 began 3 days after infection and continued until 10 (150 mg/kg) or 21 (50 mg/kg) dpi. **C)** Survival curves. Log-rank (Mantel-Cox) test with median survival. **D-F)** PBMC-associated viremia (D), nasal lavage viral titers quantified by TCID_50_ assay (E), and lymphocyte counts (F) are shown. Normal range of lymphocytes is depicted by green shading. **G)** Neutralizing antibody titers. In (D-F), symbols indicate individual animals and lines represent geometric means. Two-way ANOVA followed by Dunnet’s (comparisons to vehicle) post-hoc multiple comparisons test.

In a proof-of-concept oral efficacy study, we infected ferrets with a lethal inoculum of the wild type CDV-5804p strain and initiated treatment at 3 dpi, when PBMC-associated viremia and fever become first detectable in this model (Fig. 4B and Fig. S10A,B). Exploring two distinct treatment regimens, animals received GHP-88310 either at 50 and 150 mg/kg b.i.d., or 50 and 100 mg/kg q.d., the latter corresponding to the lowest efficacious daily dose of 100 mg/kg of GHP-88309 (*25*) and half that dose. Treatment was continued in all groups until 10 dpi (150 mg/kg b.i.d. group) or 21 dpi (50 mg/kg b.i.d. and q.d. groups). All treated animals survived the infection (Fig. 4C) and clinical signs were fully mitigated (Fig. S10A,B), whereas all animals in a vehicle-treated reference group succumbed to CDV disease with a median survival of 11 dpi. Cell-associated viremia was first detectable 3 dpi, when therapeutic treatment was initiated, and rapidly progressed in vehicle-treated animals to peak viremia titers of approximately 1.3 × 10^5^ TCID_50_ units per 10^6^ PBMCs at 7 dpi (Fig. 4D). Treatment at 50 or 100 mg/kg q.d. or 50 mg/kg b.i.d. lowered peak titers by ∼1 order of magnitude and resolved viremia with similar kinetics by 15-16 dpi. Treatment at 150 mg/kg reduced peak viremia by 2 orders of magnitude and was sterilizing at 10 dpi. The effect of treatment on shed virus load in nasal lavages closely mirrored that on viremia titers (Fig. 4E). Virus shedding from animals receiving GHP-88310 at 150 mg/kg b.i.d. peaked 2 orders of magnitude lower than in the vehicle group and shedding from treated animals ceased 6 dpi. Low b.i.d. dose animals and animals of both q.d. groups showed biphasic shedding kinetics, characterized by an initial decline after treatment start followed by a second shedding peak at approximately 10 dpi. Infectious virus was undetectable in all treated animals 17 dpi. Lymphocytopenia, a hallmark of morbillivirus disease, was efficiently mitigated by 50 mg/kg b.i.d. or 100 mg/kg q.d. GHP-88310 and fully suppressed at the 150 mg/kg b.i.d. dose level (Fig. 4F). Animals in the 50 mg/kg q.d. treatment group experienced vehicle-like lymphocytopenia until 9 dpi. Subsequently, lymphocyte counts rapidly stabilized and returned to normal range by 27 dpi. Consistent with transient lymphocytopenia, the emergence of neutralizing antibodies was delayed in animals of the 50 mg/kg q.d. cohort compared to the other treatment groups (Fig. 4G). Whole genome sequencing of treatment-experienced virus populations recovered from animals 3, 5, 8,13, and 14 dpi did not detect any mutations in polymerase proteins (Data File S1), indicating a high barrier against viral escape from GHP-88310 *in vivo*.

### Molecular interactions between GHP-88310 and the viral L protein

We have identified the central cavity of the viral P-L polymerase complex as target site for GHP-88309-class inhibitors through photo-affinity crosslinking and reduced susceptibility profiling (*23*). Several high-resolution structural models of the MeV polymerase complex were recently solved (*30, 31*), setting the stage for extraction of the GHP-88310 docking pose. Activity testing of GHP-88310 against panels of recombinant HPIV3 (Fig. 5A), recombinant SeV (Fig. 5B), and MeV minigenome (Fig. 5C) harboring mutations mediating escape from GHP-88309 demonstrated, with few exceptions, high consistency of resistance sites, suggesting conserved docking poses of both compounds. We defined substitutions that caused a >5-fold increase in EC_50_ and/or EC_90_ values as major drivers of reduced viral susceptibility (highlighted in Fig. 5A-C). Exceptions were the HPIV3 L-T1010A variant, which remained fully sensitive to GHP-88310, and SeV L-Y942H, which mediated robust resistance to GHP-88310 but moderate escape from GHP-88309. Corroborating our previous observations (*23*), residues at confirmed resistance sites were highly conserved across members of the respirovirus and morbillivirus genus (Table S2).

**Figure 5.**
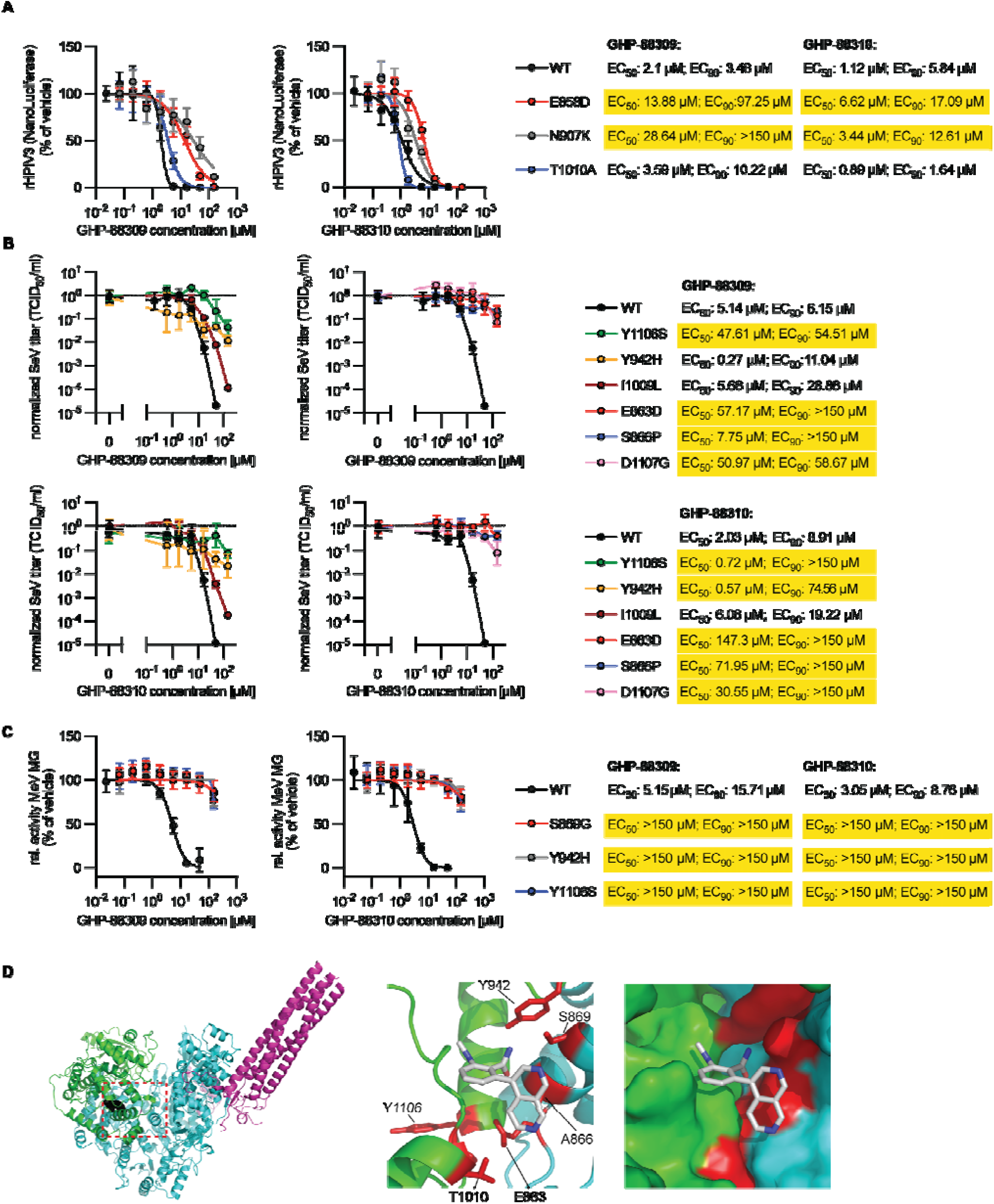
Functional and molecular profiling of GHP-88310. **A-C)** Dose-response assays with recombinant virus variants or in a minigenome harboring mutations that mediate reduced susceptibility to GHP-88309. Shown are 4-parameter variable slope non-linear regression curves for HPIV3 (A), SeV (B), and in a MeV minigenome (C). Fifty and 90% inhibitory concentrations against each compound are provided. Yellow highlights mark mutations that result in a ≥5-fold increase in EC_50_ and/or EC_90_ compared to the unmodified genetic parent virus. **D)** *In silico* docking of GHP-88310. Ribbon representation of the MeV L-P structure (PDBID: 9dus), highlighting the predicted docking location (black spheres in red dashed box) of GHP-88310 (left). The RdRP and capping domains are colored in cyan and green, respectively. Ribbon representation of the top scoring docking pose conserved between GHP-88309 and GHP-88310 (middle). GHP-88310 docking pose is shown as grey sticks. Residues that have been shown to confer resistance to GHP-88309 are highlighted in red. Surface representation of the docking site of GHP-88310 showing the location of the docking pose between the capping and RdRP domains (right). In (A-C), curves show 4-parameter variable slope non-linear regression models; n ≥3. Symbols represent means ± SD (A,C) or the geometric mean ± geometric SD (B).

Informed by these conserved mediators of resistance, photoaffinity mapping results for GHP-88309, and biolayer interferometry (BLI) binding assays of GHP-88310 to purified recombinant MeV P-L complexes (Fig. S11), we docked both compounds *in silico* into a native structural model of the MeV polymerase (PDB: 9DUS). Top scoring poses consistently placed both inhibitors into the same defined pocket at the interface between the L protein RdRP and PRNTase domains (Fig. 5D and Fig. S12). However, distinct predicted docking poses emphasized a high-affinity interaction between the GHP-88310 diazo quinoline ring and residue A866 of the RdRP domain that was not present in the GHP-88309 pose (Fig. S13A,B). Major resistance sites lined the binding pocket, predicting direct contact between the ligand and L-protein residues A866, Y942, and T1010. Additional hot-spot sites were located in close proximity, suggesting that viral escape from either inhibitor is predominantly due to primary, rather than long-range secondary, resistance.

### GHP-88310 efficacy in primary human cells and organoids

To assess whether antiviral potency of GHP-88310 supports realistic human dose concentrations, we determined efficacy in physiologically relevant primary human airway epithelium cells and well-differentiated human airway epithelium organoids. Wash-in and wash-out studies assessed intracellular exposure levels during 2-hour periods in undifferentiated human bronchial tracheal epithelial cells (HBTECs) in, respectively, the presence of 80 μM extracellular GHP-88310 (wash-in) or after removal of extracellular compound (wash-out) (Fig. S14). At wash-in, a maximal intracellular exposure plateau of approximately 10^5^ pmol/10^6^ cells was reached in 15 minutes (earliest sampling time point), indicating high plasma-membrane permeability, and was sustained for the duration of the experiment. After removal of extracellular GHP-88310, intracellular drug levels declined by approximately 2 orders of magnitude within the 2-hour monitoring window.

Efficacy testing of GHP-88310 against the primary HPIV3 indication in undifferentiated HBTECs showed sub-micromolar potency and a steep Hill slope, resulting in a fully sterilizing antiviral effect at concentrations of 10 μM and greater (Fig. 6A). After generation of well-differentiated airway epithelium organoids grown at air-liquid interface (Figure 6B), we exposed cultures to ascending concentrations of GHP-88310 in the basolateral chamber of the transwell system for 48 hours each and measured corresponding transepithelial electrical resistance (TEER) as a well-established marker of cytotoxicity in the system(*32-34*). Electrical resistance remained unchanged at approximately 600 ohm × cm^2^ before and after exposure (Fig. 6C), indicating that GHP-88310 was well-tolerated at the highest physiologically achievable concentrations without disturbing tissue integrity.

**Figure 6.**
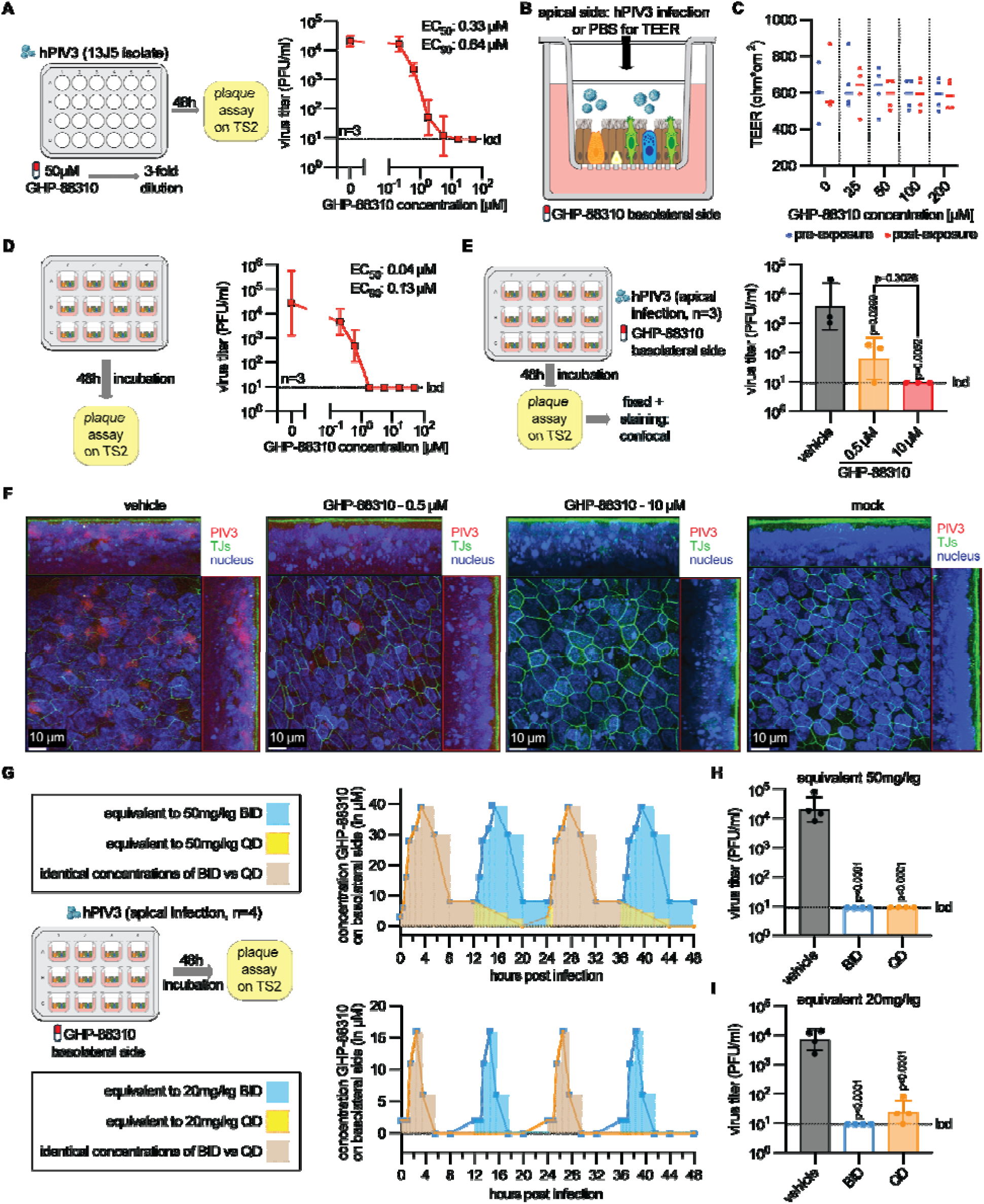
GHP-88310 efficacy in well-differentiated air liquid interface human airway epithelial cultures. **A)** Virus yield reduction of HPIV3 (clinical isolate 13J5) in undifferentiated NHBE cells (F3, n=3). **B)** Schematic of well-differentiated air-liquid interface human airway epithelial organoids. **C)** Assessment of GHP-88310 cytotoxicity in organoids. TEER was measured before and after 48 hours of exposure to increasing concentrations of GHP-88310 in the basolateral chamber. Symbols represent individual biological measurements, and lines show medians. **D)** Virus yield reduction assay of HPIV3 in organoids. Shed progeny virus was collected from the apical side of the transwells. Curves show 4-parameter variable slope non-linear regression models, symbols represent the geometric mean ± geometric SD. **E)** HPIV3 titers in infected organoids used for confocal imaging. **F)** Confocal microscopy of organoids infected with HPIV3 at 2dpi. Cells were stained with anti-HPIV3 HN (red), anti-ZO-1 (tight junctions; green), and Hoechst 34580 (nuclei; blue). Z-stacks of 100-150 0.22 µm slices with 63× oil objective. Scale bar, 10 µm. **G)** *Ex vivo* PK-guided assessment of GHP-88310 efficacy against HPIV3 in organoids. The simulation recapitulates the PK profile of GHP-88310 following 14 doses of 50 mg/kg b.i.d. (top) or the PK profile after 20 mg/kg q.d. Solid lines represent the experimentally measured PK profiles in ferrets. Corresponding compound concentrations of GHP-88310 applied at each time point are shown as bars. **H,I)** HPIV3 titers from the *ex vivo* simulation study. In (E,H,I), bars indicate the geometric mean ± geometric SD and symbols represent individual biological repeats. One-way ANOVA with Dunnet’s (comparisons to vehicle) post-hoc multiple comparisons test. Group sizes (n) are indicated in each study schematic.

Dose-response assays using static concentrations of basolateral GHP-88310 after apical infection of organoids with 5,000 plaque forming units (pfu) HPIV3 returned nanomolar EC_50_ values with favorable hill slope (EC_90_ 0.13 μM), based on titers and viral RNA copy numbers of apically released progeny virions (Fig. 6D, Fig. S15). Sterilizing conditions were reached at 1.85 μM GHP-88310 in the basolateral chamber, approximately equivalent to trough plasma concentrations in the multi-dose ferret PK studies when compound was administered at 50 mg/kg. To verify enhanced antiviral potency of HPIV3 in the organoid models compared to immortalized cell lines, we imaged HPIV3-infected, fixed, and stained organoid sections after incubation in the presence of 0.5 or 10 μM compound. Titration of progeny virions prior to imaging demonstrated that 0.5 μM basolateral GHP-88310 reduced virus titers by approximately 2 orders of magnitude (Fig. 6E), which was consistent with the dose-response assay results. Anti-ZO-1 staining revealed an unperturbed tight junction network in the presence of both concentrations examined, similar to mock-treated uninfected cultures (Fig. 6F). Only sporadic viral antigen was detectable after incubation of infected cells in the presence of 0.5 μM GHP-88310, and none after incubation with 10 μM compound, whereas HPIV3 proteins were abundant in organoids that had received vehicle volume-equivalents. Overlay of viral and beta-actin stains revealed that viral replication was concentrated in ciliated cells (Fig. S16), consistent with previous reports for HPIV3(*35, 36*).

To assess GHP-88310 efficacy in human organoids at physiological, dynamic concentrations, we recapitulated ferret plasma PK curves in the basolateral chambers of the airway epithelium organoids infected with HPIV3. Two dose levels, 20 mg/kg and 50 mg/kg, each represented by b.i.d. and q.d. dosing regimens, were simulated over a 48-hour period each (Fig. 6G). At study end, we determined progeny virus yields released from the apical surface. Validating our finding of high anti-HPIV3 potency of static GHP-88310 in the organoids, both high-dose regimens were sterilizing (Fig. 6H). Also simulating a low-dose b.i.d. treatment fully suppressed production of infectious progeny virions (Fig. 6I), although plasma PK profiles called for peak GHP-88310 concentrations of only ∼15 μM and no compound was present in the basolateral chamber for 6 hours in every 12-hour interval (Fig. 6H). Even recapitulating a 20 mg/kg q.d. plasma exposure profile in the basolateral chamber, which comprised compound-free incubation periods of 18 hours in every 24-hour interval, greatly inhibited virus replication. Progeny virus yield was statistically significantly reduced by over two orders of magnitude compared to organoid cultures that had been incubated in the presence of basolateral vehicle volume-equivalents (Fig. 6I). These results support high developmental potential of GHP-88310 for human therapy, demonstrating that potent antiviral efficacy is achieved at physiological concentrations in disease-relevant human tissue models without cytotoxicity limitations.

## Discussion

This study identified the broad-spectrum anti-orthoparamyxovirus clinical candidate GHP-88310, which showed favorable tolerability properties in rodent species and higher mammals including dogs. De-risking GHP-88309 illuminated a sharp tolerability limit in higher mammals at concentrations exceeding 100 mg/kg daily dose. We addressed this issue synthetically through development of GHP-88310, which occupies novel chemical space and was tolerated without adverse effects by ferrets and dogs at very high dose levels that far exceeded the lethal dose of GHP-88309 in ferrets. Differential responses between rodents and non-rodents to drug exposure are not unusual and often attributable to the high metabolic rate of rodents (*37-39*), an expanded rodent cytochrome P450 system that results in fast breakdown of drugs with frequently distinct metabolites (*40-42*), and/or the high liver to body weight ratio of rodents that increases resistance to chemical stress (*43*). In the case of GHP-88309, plasma exposure was high after oral dosing in both rodents and non-rodents, but only non-rodents experienced tolerability limits. At equal dose levels, GHP-88310 exposure trailed that of GHP-88309. Our ascending dose PK studies demonstrated, however, that improved tolerability of GHP-88310 was not a trivial consequence of lower exposure compared to GHP-88309, since plasma exposure in ferrets and dogs after high-dose GHP-88310 far exceeded lethal levels of GHP-88309 in ferrets. We propose that toxicity of high-dose GHP-88309 in non-rodents results most likely from accumulation of an unidentified metabolite, which is not produced in rodents due to their distinct cytochrome P450 composition, and absent from GHP-88310 breakdown products due to its distinct chemical structure.

A hallmark of viral escape from GHP-88309 is that resistance emerged readily in cell culture, but all escape variants were apathogenic and no escape mutations appeared *in vivo* (*23*). Susceptibility testing of developmental candidate GHP-88310 against recombinant strains of three viral targets – the primary indication HPIV3, surrogate respirovirus SeV, and secondary indication MeV – that each harbored confirmed GHP-88309 escape mutations (*23*) revealed overlapping susceptibility profiles. All mutations were located in the viral L protein, lining the interface between the RdRP and PRNTase domains in the central polymerase cavity. However, tolerance to GHP-88310 was, in most cases, less robust than escape from GHP-88309. Conserved susceptibility profiles spotlight a common molecular target site of both compounds, whereas reduced robustness of escape from GHP-88310 suggests altered docking poses, which was substantiated by *in silico* QSAR models. Importantly, this hallmark feature of GHP-88309 resistance appears to equally apply to GHP-88310, since whole genome sequencing of heavily treatment-experienced virus populations recovered from ferrets did not reveal substitutions at known escape hot-spots or any other polymerase residues. Although future work must address whether additional specific GHP-88310 escape sites exist, our results indicate a high barrier against viral escape from GHP compounds without concomitant loss in pathogenicity.

It was the primary focus of this synthetic lead identification program to improve tolerability in higher mammals without compromising antiviral potency, and GHP-88309 and GHP-88310 indeed have equal activity in cell culture. We therefore did not anticipate also achieving a substantial *in vivo* potency boost in both rodent and non-rodent host species. Quantitation of drug tissue levels of both compounds in cotton rats excluded improved lung tissue distribution of GHP-88310 as a possible explanation for this approximately 6-fold increase in antiviral efficacy. Whereas the lowest efficacious dose of GHP-88309 and GHP-88310 in the CDV ferret model was the same (50 mg/kg), plasma exposure of GHP-88310 was lower than that achieved by GHP-88309 after equal doses. The conclusion of an *in vivo* potency boost in both models was further supported by our observation that GHP-88310 was efficacious also in a q.d. regimen, lowering the minimally required efficacious daily dose by 50%. We propose that breakdown of GHP-88310 in both rodents and non-rodents may generate a bioactive, but non-toxic, metabolite that carries part of the antiviral efficacy, whereas GHP-88309 metabolism produces a toxic by-product. Both effects, increased tolerability and enhanced antiviral efficacy of GHP-88310, have substantially expanded the therapeutic window of the compound against the primary and secondary indication in relevant animal models, underscoring high clinical potential.

*Ex vivo*, potency of GHP-88309 against HPIV3 in established cell lines and primary HBTECs was virtually identical. However, we noted an approximately 5-fold higher antiviral potency in disease-relevant, well-differentiated human airway epithelium organoids versus the undifferentiated HBTECs. This effect may be due to a combination of the diverse cell types present in the organoids, their complex 3D-architecture, and the more natural nutrient concentration gradient compared to simple cell monolayers (*44-47*). Indeed, confocal microscopy identified, consistent with previous reports (*35, 36*), ciliated cells as the primary host cell type for HPIV3 replication in the airway epithelium cultures. To test the correlation between oral dose level and antiviral effect, we recapitulated ferret PK-informed dynamic compound concentrations in the basolateral chamber of the transwell cultures, simulating repeat oral dosing ranging from 20 mg/kg q.d. to 50 mg/kg b.i.d. We first implemented this approach when developing molnupiravir (*48*). Simulating plasma concentrations of a 20 mg/kg q.d. GHP-88310 dosing regimen in the basolateral chamber significantly reduced progeny virus yields by several orders of magnitude, which translates under consideration of established species conversion factors (*49*) to a feasible predicted oral human dose target as low as 200 mg daily. Corroborating these results, higher dose levels were fully sterilizing.

Simulation of 20 mg/kg b.i.d. or 20 mg/kg q.d. regimens entailed two 6-hour or one 18-hour window, respectively, in every 24-hour period in which no compound was present in the basolateral chamber. Effective viral inhibition despite prolonged incubation in the absence of inhibitor is likely due to inhibition of both replicase and transcriptase activity of the viral polymerase complex by GHP class compounds (*26, 50*). This concerted action may arrest the viral replication cycle in the primary transcription stage (*50*), preventing exponential expansion of viral protein expression and suppressing production of immune-modulatory viral V and C proteins that counteract the type I host interferon response (*51-54*), which exposes the virus to host cells in an unmitigated antiviral state. Both the HPIV3 cotton rat and CDV ferret models confirmed that effective virus inhibition by only intermittently present GHP-88310 was not an artifact of the *ex vivo* organoid system, since q.d. administration of GHP-88310 at the lowest efficacious dose likewise resulted in either model in prolonged periods with undetectable plasma drug levels. These results thus establish proof-of-concept that efficacious orthoparamyxovirus therapy with a non-nucleoside polymerase inhibitor does not depend on continuous drug plasma concentrations above the minimum inhibitory level, illuminating an overproportional beneficial effect of targeting the mononegavirus polymerase complex.

Naturally, human organoid systems can only partially simulate the complexity of human infection. Whereas first-in-human dose levels will be determined based on toxicology results, the organoids provide valuable insight into defining desirable dosing objectives. Other limitations of our study arise from incomplete recapitulation of all aspects of human disease presentation by animal models, currently undefined highest tolerated dose levels of GHP-88310, and limited insight into a possible effect of co-morbidities and/or low patient immune competence on antiviral efficacy. However, ferrets are a natural host for CDV and hallmarks of human morbillivirus disease are fully preserved, supporting confidence that treatment paradigms established in this model will provide valuable guidance for performance in human therapy. In summary, this study identified a much-needed novel developmental candidate for treatment of orthoparamyxovirus infections that combines attractive drug-like properties with potent antiviral efficacy and unusually broad indication spectrum, opening a viable path to clinical testing.

## Material and Methods

### Study design

Cells, normal human bronchial/tracheal epithelial (NHBE) cells, well-differentiated air liquid interface (ALI) human airway epithelial (HAE) cells, ferrets, cotton rats and mice were used as models to examine the efficacy of GHP-88310 and other GHP-88309 analogs against HPIV3, SeV, and CDV. Viruses were administered through intranasal inoculation and virus load was monitored continuously in nasal lavages (ferrets only), and in respiratory tissues of cotton rats and mice extracted when indicated after infection. Virus titers were determined through TCID_50_-titration/ plaque assays.

### Cells, viruses, and compound synthesis

Vero E6, Vero E6 TMPRSS2, Vero cells stably expressing canine signaling lymphocyte activation molecule (Vero c-SLAM, VDS), Vero-hSLAM, and BHK-T7 cells (BHK cells stably expressing T7) were cultured in Dulbecco’s modified Eagle’s medium (DMEM) supplemented with 7.5% heat-inactivated fetal bovine serum (FBS) at 37°C and 5% CO_2_. Cells were tested regularly for mycoplasma contamination prior to usage. Recombinant SeV-GFP, rCDV 5804p, HPIV3 (clinical isolate 13J5, HPIV3/Seattle/USA/13J5/2016; KY629774.1), rPIV3-nano (JS strain), rMeV (Edmonston strain), and rCedar virus (CedV)-nano were propagated on Vero E6 TMPRSS2 or VDS cells using DMEM with 7.5% serum, scraped when necessary, clarified, aliquoted, and stored at −80°C. The generation of all GHP-88309 resistant recombinant viruses used in these studies has been previously described (*23*). Virus stock titers were determined through TCID_50_-titration (rSeV, rCDV, rMeV, and rCedV) or plaque assay (PIV3), and stocks were stored in aliquots at -80°C. All virus stocks were sequence confirmed prior to usage. The general chemical synthesis strategy of GHP-88310 and other GHP-88309 analogs is depicted in Fig. S17. All newly synthesized analogs were authenticated through elemental analysis (Table S2) and stored as dry powder. NMR spectra and LC-MS traces for GHP-88310 and its analogs are shown in the Data File S1. Working aliquots of compounds were dissolved in DMSO or 1% methylcellulose for *in vitro* and *in vivo* studies, respectively.

### Primary cells

Undifferentiated cells: Normal human bronchial/tracheal epithelial cells (Lonza CC-2540S, lot number 519670, Caucasian, 42-year-old female donor “F3”, unknown cause of death, smoker) were amplified and expanded in lifeline cell culture media (LIFELINE Cell Technology) for no more than 3 passages. Cells were seeded in 24-well plates for experiments.

Differentiated ALI HAE cells (MatTek): EpiAirways (AIR-100; 23-year-old healthy male, Caucasian; grown at air-liquid interface for 13 days) were equilibrated after arrival and further differentiated in hanging-top plates (following manufacturers protocol) for 14-21 days prior to use in experiments. ALI HAE cells were fed every two days 6 ml maintenance media (MatTek), rinsed weekly and before a procedure (MatTek, TEER buffer). Transepithelial/transendothelial electrical resistance (TEER) was measured on selected inserts using an EVOM3 (World Precision Instruments) following manufacturers protocol.

### Virus titration

TCID_50_: Samples were serially diluted (10-fold starting at 1:10 initial dilution) in serum-free DMEM. Serial dilutions were added to either VDS (rCDV), Vero hSLAM (rMeV), BHKT7 (rCedV), or Vero E6 TMPRSS2 (rSeV) cells seeded in 96-well plates at 1×10^5^ cells/ml, 14 hours before infection. Infected plates were incubated for 72-96 hours at 37°C with 5% CO_2_, and titers were determined (Reed and Muench).

Plaque assays: HPIV3 virus samples were serially diluted (10-fold starting at 1:10 initial dilution) in serum-free DMEM. Serial dilutions were added to Vero E6 TMPRSS2 cells seeded in 12-well plates at 2×10^5^ cells per well, 20 hours before infection. Ninety minutes post infection, cells were overlaid with 1 ml of 2× DMEM and avicel (1:1 ratio). Infected plates were incubated for 84 hours at 37°C with 5% CO_2_. Upon incubation, cells were washed twice with PBS, stained with 1% crystal violet (in 20% EtOH), and plaques were enumerated.

### Viral RNA copy numbers

Cleared supernatants (CDV) or apical shed (HPIV3) were harvested and RNA was extracted following the manufacturers protocol (MagMAX Viral/Pathogen II Nucleic Acid Isolation Kit (applied biosystem)). RNA was examined using a Taqman Fast 1 Step (for CDV: FW: cgggcaagaaatggtcagaa, RV: ctgagcctcttccttggtga, Probe: fam 5’-acttgccgccgagcttggca-3’ bhq-1; for HPIV3: FW: tgttccagctaggacaagca, RV: ccggtcgtcacttctgtttc, Probe: fam 5’-ccacaaaccaacaggtggatcagcca-3’ bnfq) and analyzed accordingly.

### Dose-response and minigenome antiviral assays and cytotoxicity assessment

Compound stocks were prepared in dimethyl sulfoxide (DMSO) and diluted in cell culture media (final DMSO concentration of less than 0.1% in all wells). For luciferase-based dose-response assays, cells were seeded in white-walled 96-well plates one day prior to the experiment to achieve 50% confluency. Threefold serial dilutions of compounds were prepared in triplicate using an automated Nimbus liquid handler (Hamilton) and transferred to the cells. Immediately after the addition of compound, cells were infected with virus as previously described (*55*). Each plate contained four wells of positive and negative control (media containing 100 μM cycloheximide or vehicle (DMSO), respectively). Luciferase activities were determined 48 hours after infection using Nano-GLO buffer (Promega) and an H1 Synergy plate reader (Biotek). Normalized luciferase activities were analyzed with the following formula: % inhibition = (Signal_sample_ – Signal_min_)/(Signal_max_ – Signal_min_) × 100, and dose-response curves were further analyzed by normalized nonlinear regression with variable hill slope to determine EC_50_ and EC_90_.

BSR-T7/5 cells (5 × 10^4^ per well in a 96-well plate format) were transfected with plasmids encoding for P-ICB (0.01 μg), N-ICB (0.01 µg), L-ICB (0.0075 µg) and L-ICB mutant variants (0.0125 μg), and the MeV luciferase replicon reporter (0.02 μg) using GeneJuice transfection reagent. Cells were lysed 42 hours after transfection with OneGlo substrate (Promega), which was directly added to the cells, and bioluminescence was quantified using the above-mentioned formula.

Virus yield reduction assays with CDV and SeV: Cells were seeded in 24-well plates. Once reaching 90% confluency, cells were overlaid with compound dilutions (three-fold dilutions, in triplicate) and cells were infected at an MOI of 0.1. CDV: After 28 hours of incubation, the supernatant was removed, and cells were scraped in 300 µl media, subjected to two freeze-thaw cycles, clarified, and the resulting supernatant was collected for titration. SeV: Supernatant was collected after 52 hours and clarified. Samples were titrated and scored using the TCID_50_ method after 4 days.

Virus yield reduction assays in undifferentiated NHBE cells: Cells were seeded at 60% confluency in a 24-well plate. Upon reaching 80% confluency, cells were overlaid with compound dilutions (50 µM, three-fold dilutions, in triplicate) and were infected at an MOI of 0.1 with HPIV3 (clinical isolate 13J5). Supernatant was collected after 48 hours and clarified. Samples were titrated by plaque assay on Vero E6 TMPRSS2 cells and quantified after 4 days.

Virus yield reduction assays in EpiAirways: Inserts were rinsed using DPBS and infected on the apical side with 10,000 PFU of HPIV3 (clinical isolate 13J5) per well in 100 µl PBS. Sixty minutes after infection, virus was aspirated to allow for air-liquid exchange. Compound dilutions (50 µM, three-fold dilutions, in triplicate) were added on the basolateral side at the time of infection and ALI HAE cells were incubated for 48 hours (after 24 hours, the basolateral media was exchanged - same static compound concentration). After 20 hours of initial incubation, the apical surface was washed once to remove accumulated mucus. Apical shed was harvested in 300 µl PBS that had been incubated at 37°C for 30 min. Clarified samples were titrated by plaque assay on Vero E6 TMPRSS2 cells and measured after 4 days.

To assess the impact of the GHP-compound class on cell metabolic activity, Vero E6 cells were seeded at 6,000 cells per well in a 96-well plate and incubated at 37°C (5% CO_2_) following the addition of a 3-fold serial dilution of each compound, starting at 150 µM. After 72 hours of incubation, cells were incubated with 10 µl per well of PrestoBlue (ThermoFisher Scientific) for 1 hour at 37°C and fluorescence was measured using an H1 synergy plate reader (Biotek) with excitation and emission wavelengths set at 560 nm and 590 nm, respectively. Cellular metabolic activity as a surrogate for cell viability was expressed as a percentage relative to the media control.

To determine the effect of GHP-88310 on cell metabolic activity in primary cells, three to four ALI HAE cultures were exposed to increasing concentrations of GHP-88310 (25 µM - 200 µM) in maintenance media (MatTek). To assess barrier integrity, TEER was measured after 48 hours of incubation at 37°C with 5% CO_2_ and immediately prior to compound addition to the basolateral chamber of a 12-well hanging insert plate (MatTek). The measurements are displayed in Ohm × cm^2^.

### Confocal microscopy

PIV3- and mock-infected ALI HAE cells were infected and treated with GHP-88310 at either 0.5 µM or 10 µM as above. Forty-eight hours after infection the apical shed was harvested, and the cells were prepared for confocal microscopy. All incubation steps of immunostaining were performed at room temperature and in the absence of light as previously described (*48*). Briefly, the inserts were fixed for 40 min with 4% paraformaldehyde-PBS, washed once with PBS, cut, and mounted on a chamber slide (Fisher, cat# 154461PK). Cells were permeabilized with 500 µl of 0.5% Triton X-100 in PBS for 2 hours, then washed, and blocked with 500 µl per insert of 5% BSA in PBS containing 0.05% Tween-20 for 1 hour. Inserts were washed and stained with the primary antibody in an antibody-binding solution (2% BSA-PBS) for 1 hour (PIV3: goat anti-HPIV3 HN, Abcam ab28584; TJ: mouse anti-ZO-1 BD, Biosciences 610966; Mucin/Goblet cells: mouse anti-MUC5AC, ThermoFisher MA5-12175). Cells were washed three times and incubated with a conjugated antibody for 45 min (donkey anti-goat Alexa Fluor 568 ThermoFisher A-11057; goat anti-mouse IgG (H+L) highly cross-adsorbed secondary antibody, Alexa Fluor 488, Invitrogen, A-11029). Cells were washed an additional three times in PBS and then incubated for 5 min with Hoechst 34580. Following three additional washes, cells were mounted with coverslips using a drop of ProLong^TM^ Diamond Antifade Reagent (Thermo Scientific, Cat# P36970) and stored overnight.

Confocal Imaging was performed using the Zeiss Axio Observer Z.1 and an LSM 800 confocal microscope with AiryScan, controlled by the Zeiss Zen 3.1 Blue software (Windows 10). This software as well as Adobe Photoshop (25.9.0) were employed for image analysis.

### Single-dose pharmacokinetic property profiling in mice, cotton rats, ferrets, and dogs

Mice: Eight-week-old female Balb/cJ mice (The Jackson Laboratory) were acclimated for at least three days prior to study initiation. Mice were weighed and administered 20 mg/kg of GHP-class compounds by oral gavage in 1% methylcellulose (200 µl total volume). Blood samples (150 µl) were collected retro-orbitally at designated time points post dosing, centrifuged at 2,000 rpm for 5 minutes at 4°C, and the resulting plasma was stored at -80°C.

Cotton rats: Female cotton rats (Envigo), aged 2-3 months, were rested for at least three days prior to study start. Animals were weighed and dosed via oral gavage with 50 mg/kg of GHP-88309 or GHP-88310, formulated in 1% methylcellulose (500 µl total volume). Blood samples (150 µl) were collected at 30 minutes, and at 1, 2, 3, 4 and 8 hours post dosing. Plasma was processed as described above. At 8 hours post-dosing, organs (lung, brain, spleen, heart, kidney, small intestine, large intestine, and liver) were harvested and immediately frozen in liquid nitrogen.

Ferrets: Female ferrets (Triple F Farm), 4-8 months of age, were acclimated for a minimum of three days prior to dosing. Animals were weighed and administered GHP-88309 or GHP-88310 via oral gavage at doses of 150 mg/kg (GHP-88309) and 150 or 500 mg/kg (GHP-88310). Both compounds were prepared in 1% methylcellulose, with a final gavage volume of 2 ml. Blood samples (200 µl) were collected at 0.5, 1, 2, 3, 4, 6, 8, 12, and 24 hours post-dose and processed as described previously.

Dogs: Male and female beagles, aged 10-12 months old, were weighed and administered GHP-88310 orally at 50, 150, or 500 mg/kg, formulated in 1% methylcellulose (study conducted by SRI Biosciences^TM^). Blood samples (300 µl) were collected at pre-dose, 0.5, 1, 2, 4, 6, 8, 12, and 24 hours post-dosing for analysis. Clinical observations were conducted at pre-dose, immediately after dosing, 2-4 hours post-dose, and again at 24 hours. The same four animals were used across all dose levels, with a minimum washout period of seven days between each dose escalation.

For analysis, aliquots of animal plasma were mixed with 70% acetonitrile in water that included an internal standard. Samples were then vortexed for one minute and collected at 15,000 rpm for 5 minutes. The resulting supernatants were transferred to HPLC vials for analysis by LC-MS/MS at 4°C. HPLC separation was performed on an Agilent 1260 system (Agilent Technologies, Santa Clara, CA, USA) equipped with an autosampler, column oven, UV lamp, and binary pump.

For the mouse PK study screening, a Zorbax SB-Phenyl (50x4.6mm, 3.5 µm) column (Waters) was used. Mobile phase A consisted of 20 mM ammonium formate and mobile phase B consisted of acetonitrile. A 2-minute isocratic HPLC method was used at 40% mobile phase A. For the remaining PK studies, an XTerra MS C18 (50x2.1mm, 5 μm) column (Waters) was used. Mobile phase A consisted of 25 mM ammonium bicarbonate in HPLC grade water pH adjusted to 9.8 and mobile phase B consisted of acetonitrile. A 2-minute isocratic HPLC method was used at 75 or 80% mobile phase A. MS analysis was performed on a QTrap 7500 or Triple 7500 Mass Spectrometer (Sciex, Framingham, MA, USA) or using positive electrospray ionization (ESI) in multiple reaction monitoring (MRM) mode. Data analysis was performed using SciexOS Software (Sciex). PK parameters were calculated using Phoenix WinNonLin 8.4 (Build 8.4.2.6172; Certara, Princeton, NJ) using the non-compartmental analysis tool. AUC_0-8hrs_ values were calculated using GraphPad Prism (v10.1.0). Values below the limit of quantitation were plotted as LLOQ/2, following an accepted modeling strategy for PK data with individual values below detection limit (*56*).

Samples of frozen animal tissue were homogenized with 4°C 70% acetonitrile in water that included an internal standard. Homogenate were collected for 5 minutes at 10,000 rpm and supernatants subjected to a second clearance step (5 minutes, 15,000 rpm). The remaining supernatant was analyzed via LC/MS/MS, maintaining samples at 4°C. HPLC separation was performed on an Agilent 1260 system (Agilent Technologies) equipped with an autosampler, column oven, UV lamp, and binary pump, using an Xterra MS C18 (50 x 2.1 mm, 5 µm) column (Waters). Mobile phase A consisted of 25 mM ammonium bicarbonate buffer in HPLC grade water, pH 9.8, and mobile phase B consisted of acetonitrile. A 4.5-minute gradient HPLC method was performed to separate the analytes beginning with a 1-minute isocratic hold at 10% B, followed by a 2-minute gradient to 30% B, and a return to starting conditions for 1.5 minutes. MS analysis was performed on a ZenoTOF 7600 Mass Spectrometer (Sciex) using positive ESI in multiple reaction monitoring (MRM) mode. Data analysis was performed using SciexOS Software (Sciex).

### Single-dose GHP-88309 tolerability in ferrets

Female ferrets (Triple F Farm), between 6-8 months old, were acclimated for at least three days. Ferrets were weighed and dosed with GHP-88309 once via oral gavage at doses ranging from 50-1,000mg/kg. The compound was formulated in 1% methylcellulose and a final gavage volume of 2 ml was administered. Rectal temperature and clinical scores were measured before dosing and at 30 minutes, as well as 1, 2, 3, 4, 5, 6, 8, 12, and 24 hours post-dose.

Clinical scoring evaluated: i) temperature to monitor hypothermia and ii) neurological status to detect signs of acute drug toxicity. Temperature scoring was assigned as follows: 0, >38°C; 1, 37.5-37.9°C; 2, 37-37.4°C; 3, <37°C. To assess neurological status, each ferret was individually placed in a single-ventilated cage at the time of evaluation to observe its response to a novel environment. Neurological scoring was partially adapted from the Merck Veterinary Manual (*57*) and assigned as follows: 0, explores objects, weasel war dance(*58*), intact reflexes (pinch test); 1, decreased interest in exploring objects; 2, lethargic, and/or altered gait, and/or decreased reflexes; 3, obtunded, and/or seizures, and/or paralysis in one or more limbs. Upon reaching the predetermined endpoint, a clinical score of 3 in either category, ferrets were euthanized.

### Multidose tolerability studies in cotton rats and ferrets

Cotton rats: Female cotton rats (Envigo), aged 2-3 months were habituated for at least three days prior to the beginning of the study. Cotton rats were weighed and administered GHP-88310 twice daily by oral gavage at doses of 300, 600 and 1,200 mg/kg. The compound was formulated in 0.5% methylcellulose, 5 mM sodium citrate, and 0.25% Tween80, with a final gavage volume of 500 µl. Each animal received a total of 14 doses and was monitored daily for bodyweight and temperature using implantable electronic ID transponders (TP-1000, BMDS). Following the final dose, blood samples (150 µl) were collected at designated time points, centrifuged at 2,000 rpm for 5 minutes at 4°C, and the plasma was stored at -80°C.

Ferrets: Female ferrets (Triple F Farm), between 6-8 months old, were allowed to rest for at least three days. Ferrets were weighed and dosed with GHP-88309 or GHP-88310 twice daily via oral gavage at doses ranging from 50-1,000 mg/kg. The compounds were formulated in 1% methylcellulose, with a final gavage volume of 2 ml. Animals were weighed daily, and body temperature was measured using a rectal probe. Ferrets received a total of 14 doses under a twice-daily dosing regimen or 7 doses under a once-daily regimen. Blood samples were collected on days 0, 2, 4, 5, 6 and 7 post-dosing for complete blood cell (CBC) analysis, and on days 1 or 2 and 7 for blood chemistry analysis. CBC analysis was performed using a Vetscan HM5 hematology analyzer (Abaxis), following the manufacturer’s protocol. For the 500 mg/kg and 1,000 mg/kg dosing groups, as well as the multi-dose once-daily dosing study, maximum plasma concentration (C_max_) at 5 hours post-dose was measured after doses 1, 2, 4, 6, 8, 10, 12, and 14, or daily (once-daily regimen only). Blood samples (150 µl) were collected for plasma preparation as previously described. Tissues were harvested 12 hours post dosing in the 50 and 150 mg/kg groups (GHP-88310 only) and flash frozen before analyzed.

### *In vivo* infections with SeV, HPIV3 and CDV

Efficacy studies in mice: Female 129x1/SvJ mice (The Jackson Laboratory), 6 weeks of age, were acclimated for at least three days before being randomly assigned to study groups and housed under ABSL-2 conditions for infections with sequence-validated rSeV. Bodyweight and rectal temperature were measured once daily. For infection, mice were anesthetized with isoflurane and intranasally inoculated with 5×10^6^ TCID_50_ units/ml of rSeV (*23*) in a total volume of 50 µl PBS (25 µl per nare). Treatment was initiated 24 hours post-infection and administered twice-daily via oral gavage at a dose of 150 mg/kg in 200 µl of 1% methylcellulose, continuing through study day 8. A subset of animals was euthanized either five days post-infection or upon reaching a predetermined endpoint, and tracheas and lungs were collected for analysis.

Efficacy studies in cotton rats: Female cotton rats (Envigo), 3-4 weeks of age, were allowed to rest for at least three days prior to being randomly assigned to study groups and single-housed under ABSL-2 conditions for infections with sequence-validated HPIV3 clinical isolate 13J5. Bodyweight and body temperature, using an implanted electronic ID transponder (TP-1000, BMDS), were measured once daily. For infection, cotton rats were anesthetized with isoflurane and intranasally inoculated with 1×10^7^ pfu/ml of HPIV3 in a total volume of 100 µl PBS (50 µl per nare). For therapeutic dosing, animals were anesthetized with isoflurane 12 hours post-infection and administered GHP-88309 or GHP-88310 in 500 µl of 1% methylcellulose via oral gavage, either once or twice-daily, through study day 3.

Efficacy studies in ferrets: CDV unvaccinated female ferrets (Triple F Farms), 8 months of age, were rested for one week prior to random assignment into study groups and housed in groups of three animals under ABSL-2 condition. Bodyweight and rectal temperature were recorded once daily. Ferrets were anesthetized with dexmedetomidine/ketamine and intranasally inoculated with 2×10^5^ TCID_50_ units of CDV 5804p in a total volume of 200 µl PBS (100 µl per nare) (*25*). Daily, nasal lavages were performed using 1 ml of PBS supplemented with 2× antibiotics-antimycotics (Gibco). Peripheral blood mononuclear cells (PBMCs) were collected twice weekly for titration of cell-associated viremia and CBC analysis. On days 17, 21, and 35 post-infection blood samples were collected to assess neutralizing antibody titers. For therapeutic treatments, ferrets were administered GHP-88310 by oral gavage following either a once or twice-daily regimen (50-300 mg/kg) in 2 ml of 1% methylcellulose, beginning at 3-days post-infection. Ferrets were euthanized either upon reaching a predetermined study endpoint or at the conclusion of the study on day 35 post-infection.

### Virus titration of tissue samples (mice and cotton rats) and nasal lavages (ferrets only)

Organs were weighed and homogenized in 300 µl PBS using a Bead Blaster 24R (Benchmark) set to three 30 second cycles at 4°C, with one minute rest intervals between cycles. Homogenates and ferret lavage samples were clarified by centrifugation at 13,000 × g for 10 minutes at 4°C, then subsequently titrated. Viral titers were expressed as TCID_50_ units/plaques per gram input tissue or per ml.

### Virus titration of PBMCs (ferrets only)

Blood samples were collected at specified time points twice per week. As previously described (*25*), viremia was assessed by isolating PBMCs from 1 ml of whole blood. Red blood cells were lysed using ACK buffer (150 mM NH_4_Cl, 10 mM KHCO_3_, 10 µM EDTA pH 7.4) for 7 minutes, followed by two PBS washes and resuspension in DMEM. PBMC titers were determined by co-culturing serial dilutions of purified PBMCs with VDS cells and expressed as TCID_50_ per 10^6^ PBMCs.

### Neutralizing antibody assay (ferrets only)

Blood samples (300 µl) were collected on days 17 and 21 post-CDV infection (and on day 35 for the ferret CDV QD study only) to obtain plasma, which was subjected to heat inactivation (56°C, 30 minutes) and clearance centrifugation (4,000 × g, 5 minutes). Heat-inactivated and clarified plasma was diluted two-fold in serum free DMEM, with a final volume of 50 µl per well. Fifty microliters of rCDV 5804p (2×10^3^ TCID_50_ units) was added to the plasma dilutions, followed by incubation of the plasma-virus mixture at 37°C for 75 minutes. Subsequently, the mixture was added in duplicate to VDS cells in 96-well plates, incubated for 3 days, and cytopathic effect was assessed by light microscopy.

### Whole genome sequencing of virus stocks and ferret nasal lavage samples

Whole genome sequencing (WGS) was performed using metagenomic next-generation sequencing (mNGS) as described previously (*59*). Briefly, extracted RNA was treated with a TURBO DNA-free Kit (Thermo Fisher, product #AM1907) to remove DNA. RNA was then reverse transcribed using random hexamers (Thermo Fisher, product #N8080127) and Superscript IV (Thermo Fisher, product #18090010), following the manufacturer’s protocol. Double-stranded cDNA synthesis was performed using Sequenase v2.0 (Thermo Fisher, product #70775Z1000UN), and the resulting cDNA was purified using 1.8× AMPure XP magnetic beads (Beckman Coulter, product # A63882).

Metagenomic viral WGS libraries were created from purified double-stranded cDNA with tagmentation reagents from the Illumina DNA Prep with Enrichment Kit (Illumina, product #20025524), followed by 14 cycles of dual-indexed PCR for SeV and 18 cycles for HPIV3 and CDV isolates. Amplified libraries were cleaned with 0.8× AMPure XP magnetic beads. Viral isolate libraries were not subjected to enrichment and proceeded directly to sequencing. Libraries were sequenced 2×150 bp on NextSeq 2000 (CDV, HPIV3) or NovaSeq 6000 (SeV).

Due to its high viral titer, the sample “CDV ferret study - BID treatment – 50 mg/kg - ferret 2 – D8pI” was sequenced using metagenomic WGS on NextSeq 2000 with 1×100 bp read format.

Viral WGS was also performed on all ferret nasal lavage samples with detectable PFU titers, except “CDV ferret study - BID treatment – 50 mg/kg - ferret 2 – D8pI”. These samples were processed using QIAseq xHYB Microbial Hyb&Lib Kit A (Qiagen, product #334525) and a custom Qiagen hybridization panel containing probes designed from CDV strain 5804 (AY386315.1) containing eGFP reporter (Qiagen, product # 334586). Following the manufacturer’s protocol, extracted RNA was depleted of ribosomal RNA and converted to double-stranded cDNA. Double-stranded cDNA was then enzymatically fragmented, end-repaired, indexed, purified with 0.9× and 1.1× bead cleanup, and amplified with 14 cycles of PCR, followed by a final bead purification, according to the manufacturer’s protocol.

Pre-capture sequencing libraries were pooled based on PFU titer, with one to four samples per pool, and incubated with the custom capture panel. After overnight hybridization with biotinylated probes, probe-target hybrids were bound to streptavidin-coated magnetic beads and washed to remove unbound library fragments. The enriched libraries were amplified with 20 cycles of post-hybridization PCR and purified via 1.1× bead clean-up. Post-amplification library fragment sizes were estimated with the TapeStation 4200 D1000 (Agilent), and concentrations were measured with Qubit 4 Fluorometer and Qubit dsDNA HS Assay Kit. Libraries were sequenced on Illumina NextSeq 2000 instruments using a 1×100 bp read format.

Data analysis: Raw reads were trimmed and quality filtered using fastp (v0.23.4)(*60*) with the following parameters: --cut_mean_quality 20 --cut_front --cut_tail --length_required 20 --low_complexity_filter --trim_poly_g --trim_poly_x. The consensus sequence of “recCDV_5804p WT” was generated by majority voting with Geneious Prime 2025.0, using the CDV plasmid sequence (AY386315.1 background) containing eGFP reporter as a reference.

Variants were identified using RAVA workflow with default parameters (https://github.com/greninger-lab/RAVA_Pipeline/tree/publication_GSU_2025_03_01)(*61*), (*62*). References used in RAVA analysis: for “recCDV_5804p WT” we used plasmid map as a reference (AY386315.1 background), for the “HPIV3 clinical isolate 13J5” – HPIV3/Seattle/USA/13J5/2016 (KY629774.1), for “recSeV_GFP” – Respirovirus muris strain 52 (MH557085.1), and, finally, for CDV ferret nasal lavage samples – “recCDV_5804p WT” consensus sequence. Sample “CDV ferret study - BID treatment – 50 mg/kg - ferret 2 – D8pI” had insufficient coverage (mean coverage of ∼49x with some CDS regions having 0x coverage) and was therefore excluded from the RAVA analysis. Complete lists of identified non-synonymous and complex SNVs (allelic frequency ≥ 5%, sequencing depth of the nucleotide position ≥ 30) can be found in the Data File S2. Data availability: raw sequencing data is publicly available in NCBI BioProject PRJNA1217449.

### Preparation of recombinant MeV P-L complexes

MeV P and L complexes were expressed in SF9 cells using the pFastBac Dual (ThermoFisher Scientific) expression system as previously described (*23, 63*). For purification, cells were lysed 76 hours after infection in a 20 mM imidazole, 50 mM NaH_2_PO_4_, pH 7.5, 150 mM NaCl, and 0.5% NP-40 buffer and purified by Ni-NTA affinity chromatography. L-P complexes were eluted in 250 mM imidazole, 50 mM NaH_2_PO_4_, pH 7.5, 150 mM NaCl, and 0.5% NP-40, followed by buffer exchange to 150 mM NaCl, 20 mM Tris-HCl, pH 7.4, 1 mM DTT and 10% glycerol using Zeba spin columns. Protein preparations were aliquoted and stored at -80°C until use.

### BLI analysis of GHP-88310 binding to purified MeV P-L complexes

BLI-based dissociation kinetics of GHP-88310 were determined as previously described (*23, 63, 64*). Briefly, purified MeV L-P complexes were buffer-exchanged to PBS (pH 7.4; room temperature) using Zeba spin desalting columns (7K MWCO, 0.5 ml, ThermoFisher Scientific) and biotinylated with EZ-Link Sulfo-NHS-SS-Biotin reagent (ThermoFisher Scientific). Biotinylation reactions were stopped after 30 minutes and buffer exchanged to PBS using Zeba spin columns, followed by loading on Super-Streptavidin sensors (Sartorius) for 30 hours at 30°C to reach a shift of ≥1 nm. Uncoupled streptavidin was quenched for 5 min with 2 mM biocytin solution. A solution of thyroglobulin (1 mg/ml; GE Healthcare) was biotinylated in parallel and loaded to ≥1-nm shift on reference sensors. Kinetic experiments were performed at 30°C with 800 rpm shaking in 96-well plates using an Octet Red 96 system (ForteBio). Biosensors loaded with L-P complexes or thyroglobulin were equilibrated for 120 seconds in kinetic assay buffer (0.01% BSA, 0.005% Tween 20, and 1% DMSO in PBS) to establish baselines and subsequently incubated with ascending concentrations of GHP-88310 (3-fold increases from 0.96 µM to 26 μM) for 150 seconds and association recorded. After each incubation step with GHP-88310, sensors were intermittently incubated in assay buffer for 150 seconds and dissociation recorded. Real-time dissociation kinetics (K_D_) were determined using the Octet Red software package (ForteBio Data Analysis Software, 8.2.0.7). Raw signals were processed using the double-reference method, subtracting both the non-specific thyroglobulin signal and the drift signal obtained in the absence of compound after association alignment and interstep correction at the dissociation. Binding constants were extracted from association and dissociation signals using global fitting with a 1:1 model.

### In silico docking

Docking studies were performed with MOE (version 2024.06). A structure of MeV L (Protein Data Bank (PDB) 9dus) was used for the docking studies. After protonation and energy minimization, the triangle matcher method was used for initial compound placement, followed by refinement using an induced-fit protocol to dock GHP-88310 or GHP-88309 in the L structure based on resistance data information. For MeV L, residues Y942, A866, and I1009 were selected to identify the target site for docking and resulting docking poses from each polymerase target were compared. The top scoring docking poses conserved between GHP-88309 and GHP-88310 were chosen for further analysis.

### Cellular uptake and clearance in undifferentiated NHBE cells

Primary normal HBTECs (donor “F3”) were seeded in 48 well plates and utilized for a wash-in/wash-out study once confluency was achieved.

Wash-in study: An 80 µM dilution of GHP-88310 was prepared in pre-warmed Lifeline media. Following media removal from the cells, 0.5 ml of either the 80 µM compound or vehicle (volume equivalent DMSO)-media mixture was added to the wells. Plates were incubated at 37°C, 5% CO_2_ and extracted in triplicate at the indicated time points. Cells were washed twice with cold DPBS, followed by the addition of 250 µl of 70% methanol/30% water spiked with the internal standard to each well treated with GHP-88310. A mock-treated blank plate (16 well) was extracted with 250 µl of 70% methanol/30% water (no internal standard) per well at one-hour post-dosing. All samples were mixed by pipetting and transferred to 1.5 ml microcentrifuge tubes. After centrifugation at 16,000 × g for 10 minutes at 4°C, the supernatants were transferred to 2 ml tubes and stored at -80°C until LC/MS/MS analysis.

Wash-out study: An 80 µM dilution of GHP-88310 was prepared in pre-warmed lifeline media. After media removal, 0.5 ml of the 80 µM compound-media mixture was added to the wells. Plates were incubated at 37°C, 5% CO_2_ for one hour, washed twice with 0.5 ml warm DPBS, and compound free pre-warmed Lifeline media was added. Plates were further incubated as specified, followed by washing (twice with 0.5 ml cold DPBS) and extraction, processing, and analysis as described above.

### Recapitulation of ferret plasma PK profile in EpiAirways

Air liquid interface HAE cells were exposed basolaterally to dynamic GHP-88310 concentrations reflected ferret plasma levels after either 50 mg/kg oral twice-daily dosing on day 7 or a single oral dose of 20 mg/kg. All ALI HAE cell cultures (n=4 per condition: vehicle, 50 or 20 mg/kg bid, and 50 or 20 mg/kg qd) were infected with 1×10^4^ PFU per well of the HPIV3 clinical isolate 13J5 one hour prior to treatment. Infected cultures underwent either four 12-hour treatment cycles (bid) or two 24-hour treatment cycles (qd). Twenty hours post infection the apical side was rinsed once with 400 µl PBS to reduce mucus accumulation. After 48 hours post-infection, apically shed virus was collected by incubating the cultures with 300 µl of DPBS at 37°C for 30 minutes. Viral titers were then determined by plaque assay on Vero E6 TMPRSS2 cells.

### Statistical analysis and data preparation

For statistical analysis of studies comparing more than two study groups, a one-way analysis of variance (ANOVA) or two-way ANOVA with multiple comparison *post hoc* tests as specified were used to assess statistical difference between samples. An unpaired, two-tailed t-test was used to compare compound concentrations across the analyzed organs. Statistical analyses were performed using GraphPad Prism (v10.1.0). All figures were assembled in GraphPad Prism and Adobe Illustrator/Photoshop. Study schematics were created using Adobe Illustrator, with animal illustrations generated via the vector tool. Objects in figures 5 and 6 were drawn in Adobe Illustrator and adapted from BioRender. The number of individual biological replicates (n values) and exact P values are shown in the figures. The threshold of statistical significance (α) was set to 0.05. Source data and statistical analysis are shown in the source data (Data File S3) and statistical analysis (Data File S4) files, respectively.

### Ethical compliance

All animal work was performed in compliance with the *Guide for the Care and Use of Laboratory Animals* of the National Institutes of Health and the Animal Welfare Act Code of Federal Regulations. Experiments involving mice, cotton rats and ferrets were approved by the Georgia State University Institutional Animal Care and Use Committee (IACUC) under protocols A23011, A24022 and A22035, respectively. All experiments using infectious material were approved by the Georgia State University Institutional Biosafety Committee (IBCs) and performed in BSL-2/ABSL-2 containment facilities, respectively.

## Supporting information

Lieber et al_Supplementary Information_rev

## Materials & Correspondence

Correspondence and material requests should be addressed to R.K. Plemper

## Acknowledgements

We thank the Georgia State University Department of Animal Resources for expert assistance and M. Sirrine for assistance. The recHPIV3 and recCDV-5804p genomic plasmids and Vero-canine SLAM cells were kind gifts of B. Lee, V. von Messling, and Y. Yanagi, respectively. Blood chemistry analysis was provided by the Pathology Core of Emory University.

## Author Contributions

Conceptualization: MGN, RKP

Methodology: CML, ZMS, ALG, MGN, RKP

Investigation: CML, JDW, MG, JJY, ZMS, CER, AIL, LAH, DV, AAC, MKA, REK, RMC

Visualization: CML, JDW, ALG

Supervision: GRP, ALG, MGN, RKP

Writing – original draft: CML, RKP

Writing – review & editing: CML, ZMS, GRP, ALG, RKP

## Data and Materials Availability

All data needed to evaluate the conclusions in the paper are present in the paper, the Supplementary Materials, and/or the accompanying numerical source dataset Data File S3.

## Table of Contents of Supplementary Materials

Figure S1. Multi-dose tolerability of GHP-88309 in ferrets

Figure S2. Clinical parameters of SeV infection in mice

Figure S3. Clinical parameters and viral titers of cotton rats infected with HPIV3

Figure S4. Multi-dose tolerability in cotton rats

Figure S5. Once daily administration of GHP-88310 against HPIV3 in cotton rats

Figure S6. Multi-dose tolerability and CBC parameters in ferrets

Figure S7. Ferret tissue distribution of GHP-88310 12 hours post dosing

Figure S8. Clinical parameters of multi-dose GHP-88309 and GHP-88310 tolerability

Figure S9. Quantitation of CDV RNA copy numbers through qRT-PCR

Figure S10. Evaluation of once- and twice-daily GHP-88310 dosing for CDV infection in ferrets

Figure S11. *In vitro* binding of GHP-88310 to the viral polymerase

Figure S12. *In silico* docking of GHP-88309 Figure S13 2D docking poses

Figure S14. Cellular uptake and retention of GHP-88310

Figure S15. Quantitation of HPIV3 RNA copy numbers through qRT-PCR

Figure S16. Confocal imaging of GHP-88310 antiviral activity in human airway epithelium organoids

Figure S17. Chemical synthesis strategy of GHP-88310 and its analogs

Table S1. Elemental analysis of GHP-88309 analogs

Table S2. GHP-88310-class resistance sites

