## Supplementary material for "Developmental candidate GHP-88310/EIDD-3608 with high tolerability and oral efficacy in measles and respiratory paramyxovirus models": Lieber et al_Supplementary Information_rev

**Supplementary figures**

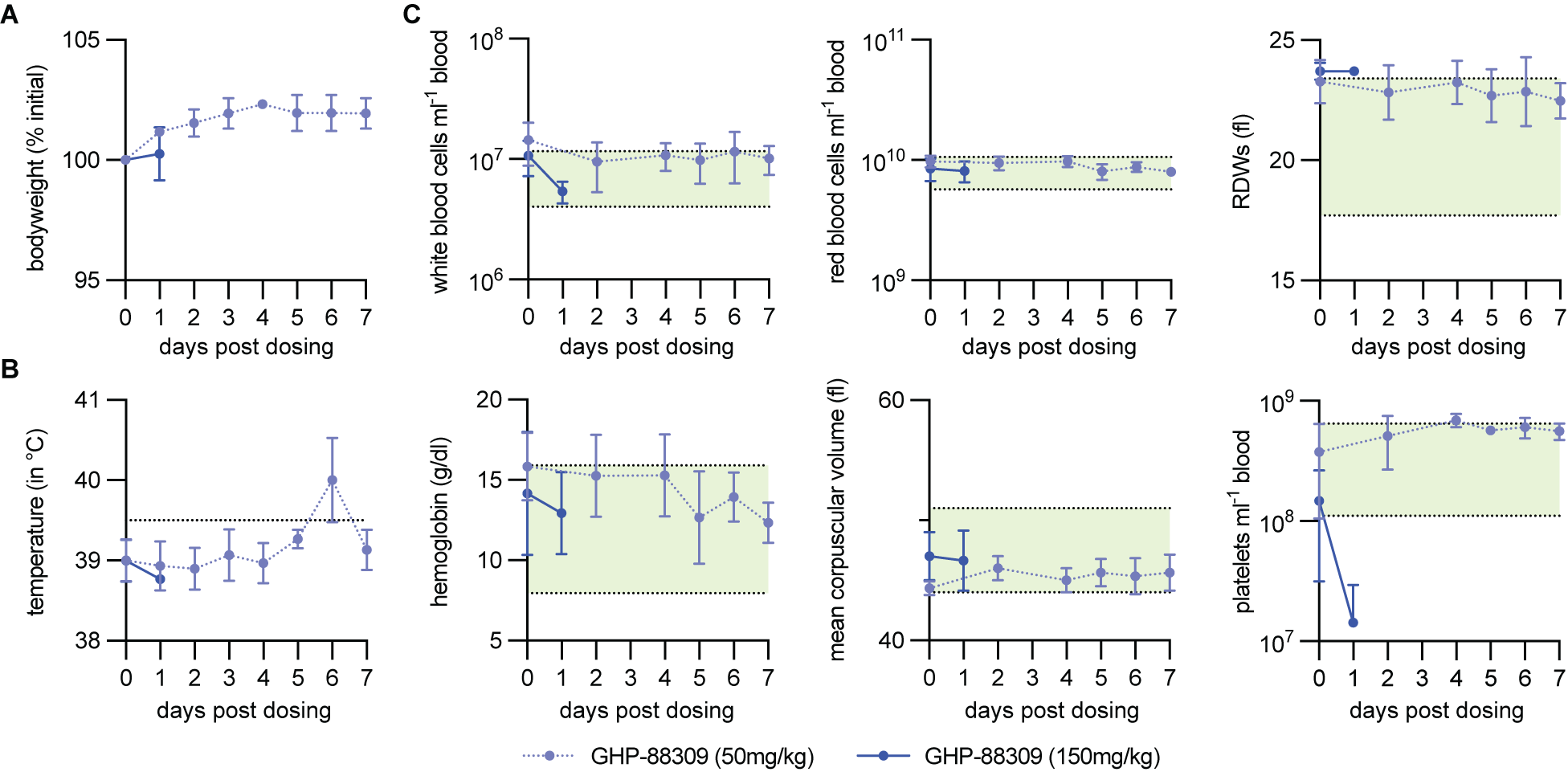

**Figure S1. Multi-dose tolerability of GHP-88309 in ferrets. A,B)** Bodyweight (A) and body temperature (B). Fever (≥39.5°C) is indicated by a horizontal dashed line. **C)** CBC analysis of ferret blood samples. Normal range for each parameter is shown in green shading. In all panels, symbols represent means ± SD.

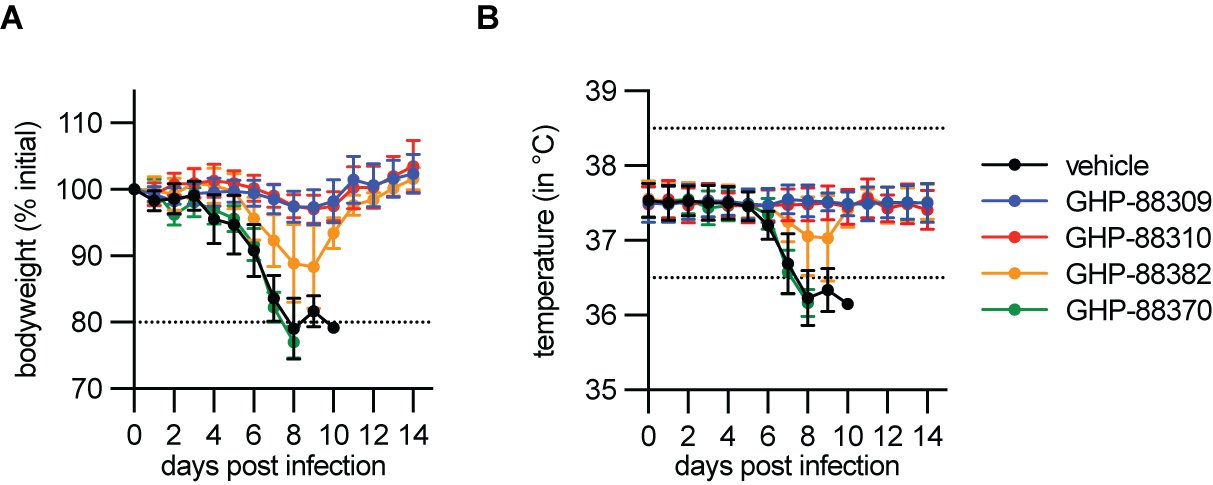

**Figure S2. Clinical parameters of SeV infection in mice. A)** Bodyweight. Predefined weight loss endpoint (80%) is shown as a horizontal dashed line. **B)** Body temperature. Normal temperature range (36.5 – 38.5°C) is shown as horizontal dashed line. In both panels, symbols represent means ± SD.

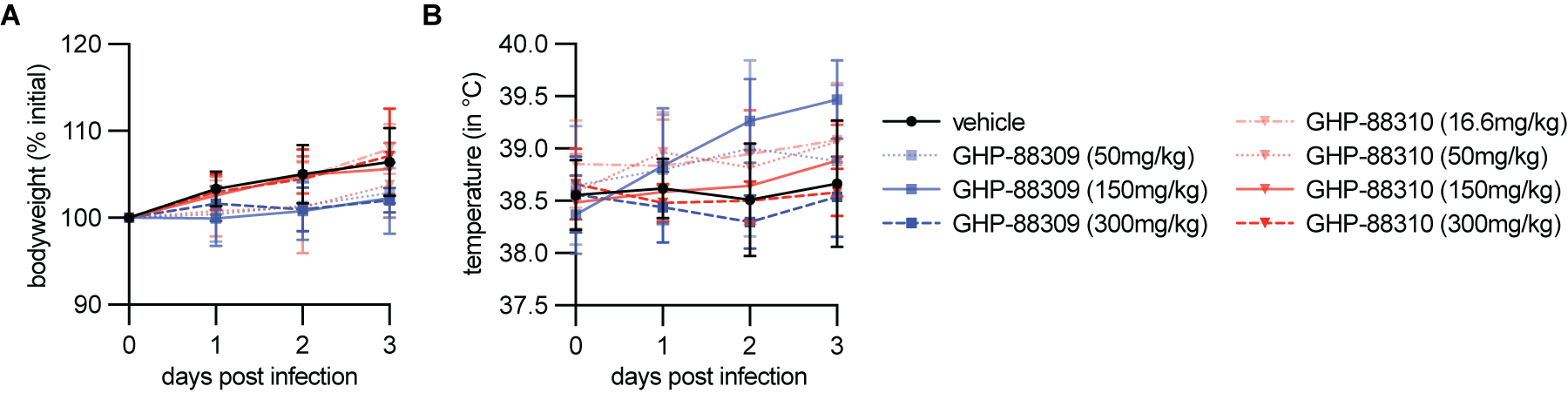

**Figure S3. Clinical parameters and viral titers of cotton rats infected with HPIV3. A,B)** Bodyweight (A) and body temperature (B) of HPIV3-infected animals. Symbols represent means ± SD.

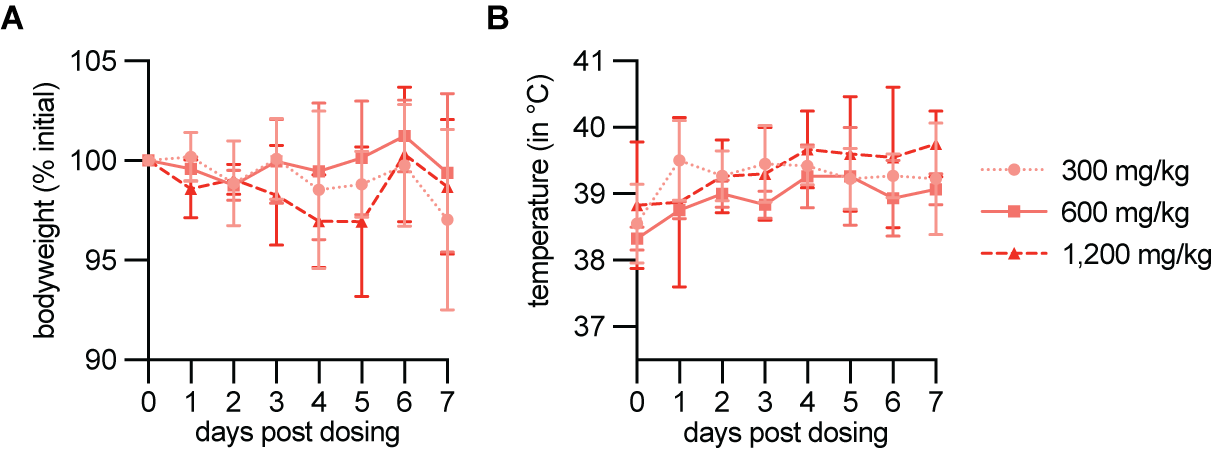

**Figure S4. Multi-dose tolerability in cotton rats. A,B)** Bodyweight (A) and body temperature (B). Symbols represent means ± SD.

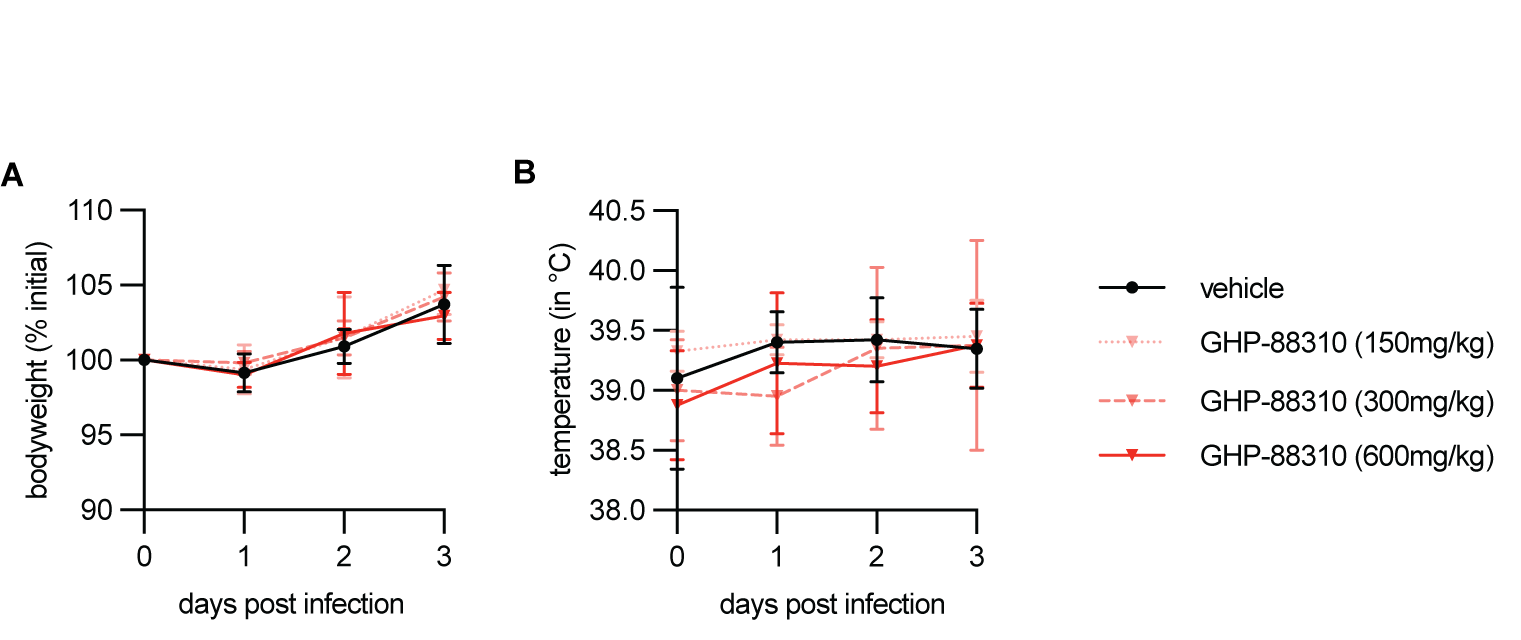

**Figure S5. Once daily administration of GHP-88310 against HPIV3 in cotton rats. A,B)** Bodyweight (A) and body temperature (B) of HPIV3-infected animals. Symbols represent means ± SD.

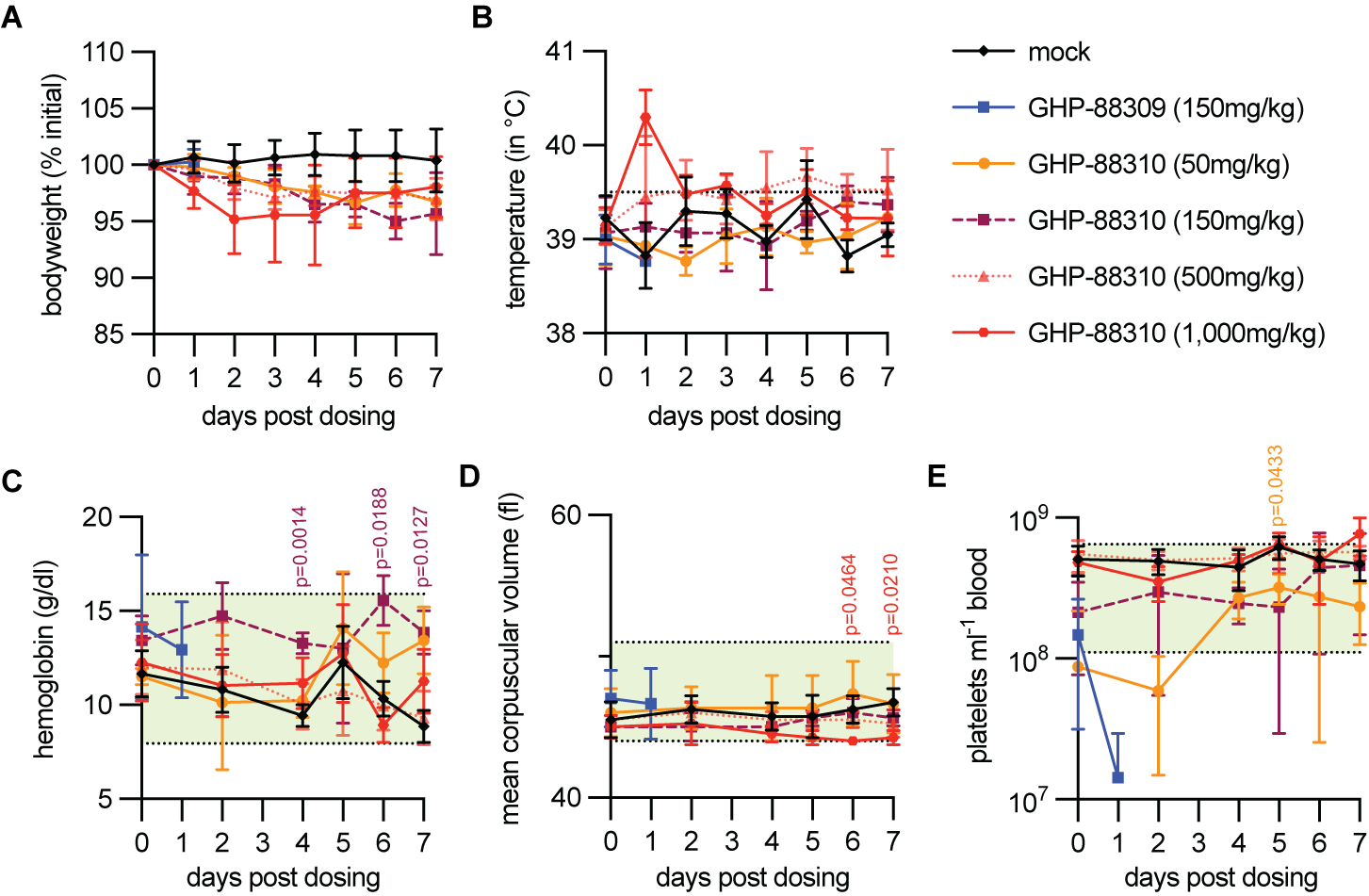

**Figure S6. Multi-dose tolerability and CBC parameters in ferrets. A,B)** Bodyweight (A) and body temperature (B). Fever (≥39.5°C) indicated by a horizontal dashed line. **C-E)** CBC analysis of ferret blood samples. Normal range for each parameter is indicated in green shading. In all panels, symbols represent means ± SD. Two-way ANOVA followed by Dunnet’s (comparisons to mock) post-hoc multiple comparisons test.

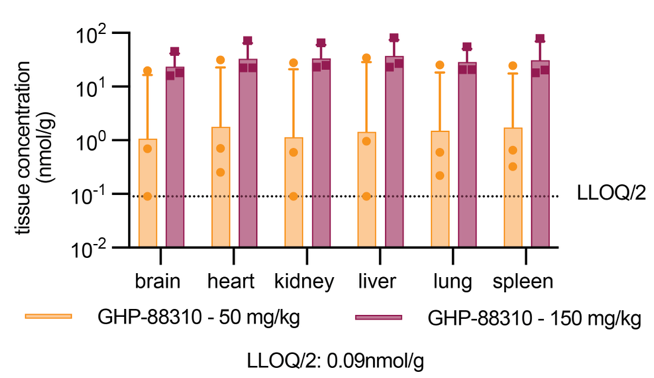

**Figure S7. Ferret tissue distribution of GHP-88310 12 hours after dosing.** Soft organ concentrations of GHP-88310 were determined 12 hours after the last of 14 doses, administered in a b.i.d. regimen. Columns indicate geometric means ± geometric SD, symbols denote individual animals. LLOQ, lower limit of quantitation.

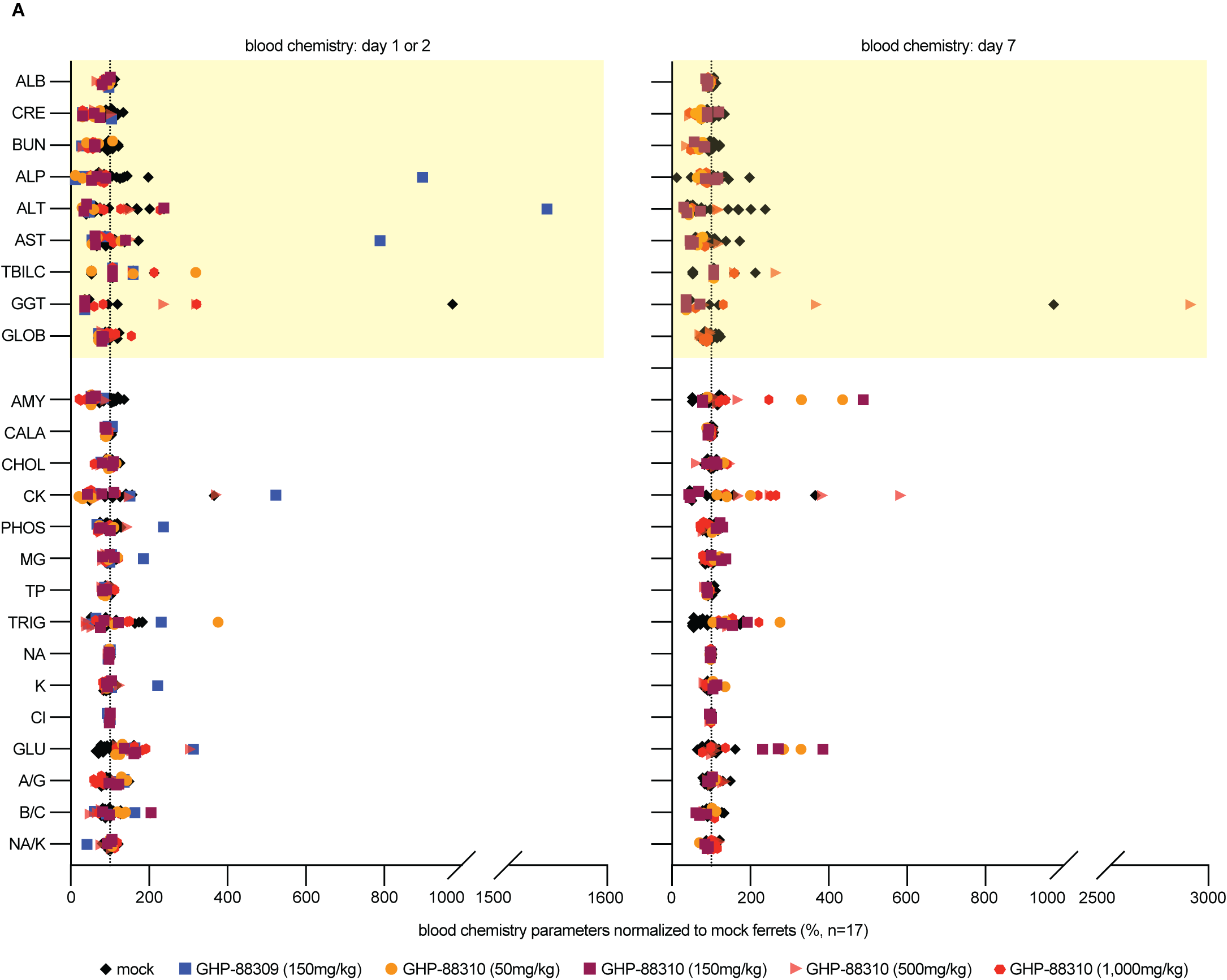

**Figure S8. Clinical parameters** **of multi-dose GHP-88309 and GHP-88310 tolerability. A)** Blood chemistry analysis of serum collected on days 1 or 2, and day 7 after dosing. Serum from the 150 mg/kg GHP-88309 group was collected on day 1 only. Key kidney and liver parameters are highlighted in yellow shading. Values were normalized to those of mock ferrets (n=17) and are presented as percent increase or decrease. Dashed vertical line indicates unchanged to mock-dosed animals (100% mark). Symbols represent individual animals. ALB, albumin; CRE, creatine; BUN, blood urea nitrogen; ALP, alkaline phosphatase; ALT, alanine transaminase; AST, aspartate aminotransferase; TBILC, total bilirubin; GGT, gamma-glutamyl transferase; GLOB, globulins; AMY, amylase; CALA, calcium; CHOL, cholesterol; CK, creatine kinase; PHOS, phosphate; MG, magnesium; TP, total protein; TRIG, triglycerides; NA, sodium; K, potassium; CL, chloride; GLU, glucose; A/G, ratio of albumin to globulin; B/C, ratio of BUN to creatine; NA/K, ratio of sodium to potassium.

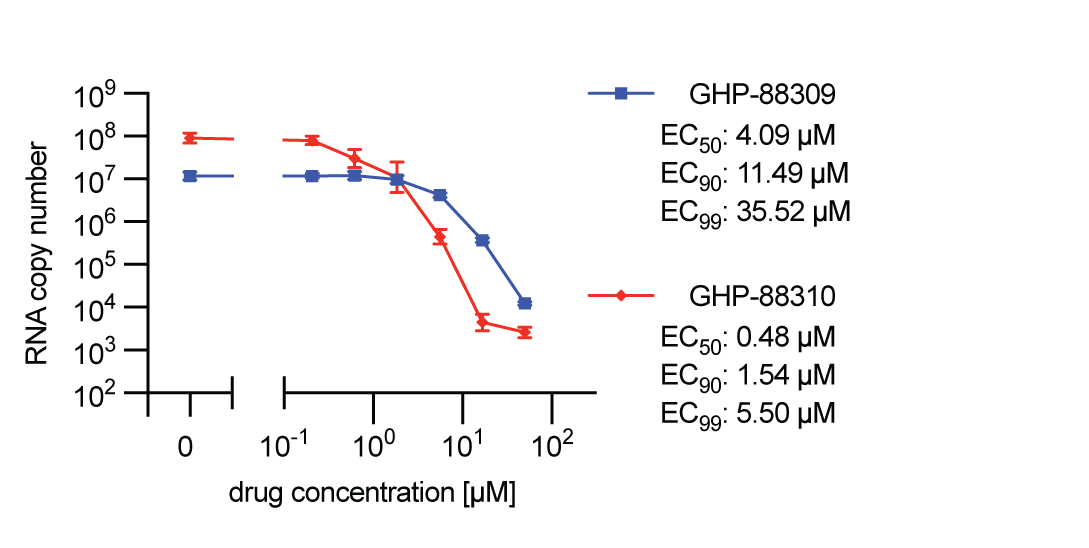

**Figure S9. Quantitation of CDV RNA copy numbers through qRT-PCR.** Viral RNA copy numbers detected in samples of the dose-response assay shown in Fig. 4A. Active concentrations were calculated based on 4-parameter variable slope non-linear regression models. Symbols indicate geometric means ± geometric SD, lines connect geometric means; n ≥3.

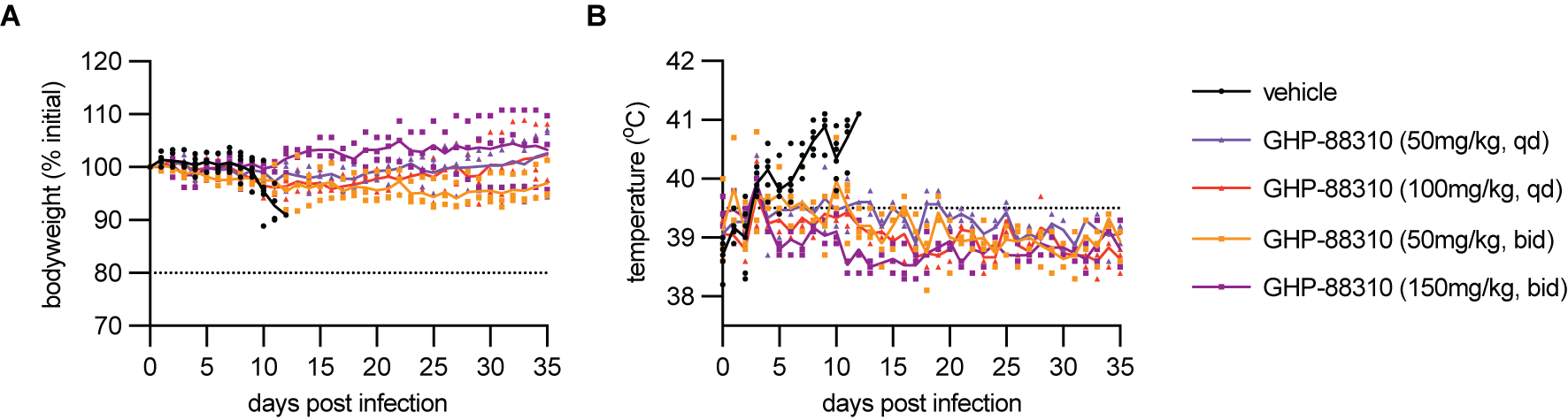

**Figure S10. Evaluation of once- and twice-daily GHP-88310 dosing for CDV infection in ferrets. A,B)** Bodyweight (A) and body temperature (B). Predefined weight loss endpoint (80%) and fever (≥39.5°C) are shown by horizontal dashed lines. In both panels, symbols represent means ± SD.

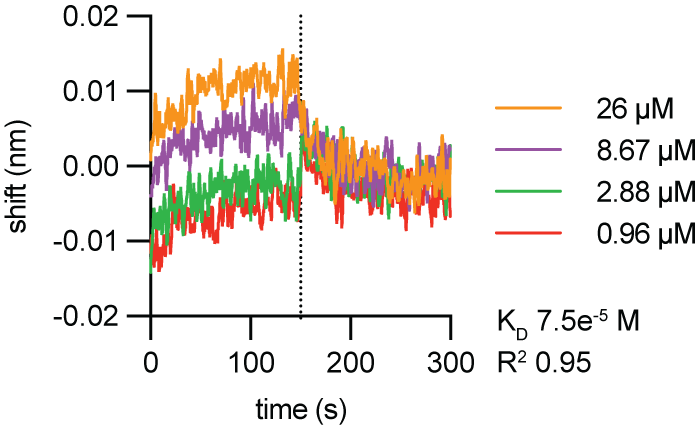

**Figure S11. *In vitro* binding of GHP-88310 to the viral polymerase.** BLI of GHP-88310 and purified recombinant MeV P-L complexes, testing different compound concentrations. Dashed vertical line denotes transition from association to dissociation. Calculated dissociation constant (K_D_) and goodness of fit (R^2^) are shown.

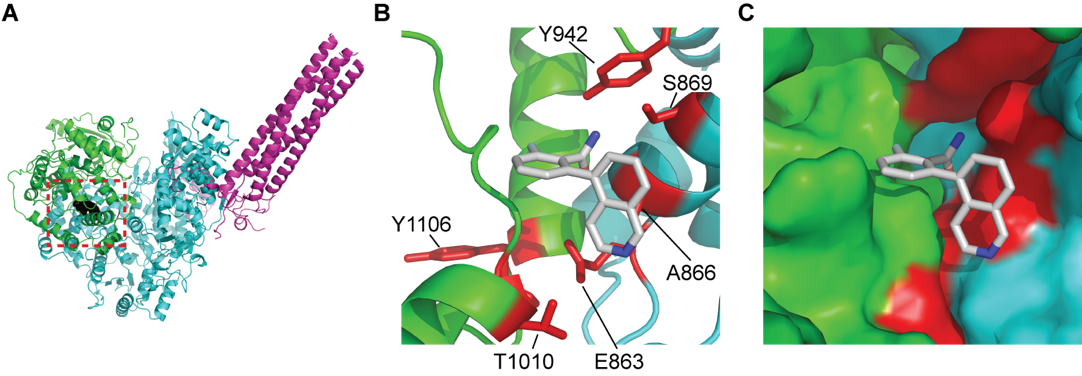

**Figure S12. *In silico* docking of GHP-88309.** **A)** Ribbon representation of the MeV L-P structure (PDBID: 9dus) highlighting the docking location of GHP-88309 (black spheres in red dashed box). The RdRP and capping domains are colored in cyan and green, respectively. **B)** Ribbon representation of the top scoring docking pose conserved between GHP-88309 and GHP-88310. GHP-88309 docking pose is shown as grey sticks. Residues that have been shown to confer resistance to GHP-88309 are shown in red. **C)** Surface representation of the docking site of GHP-88309 showing the location of the predicted docking pose between the capping and RdRP domains.

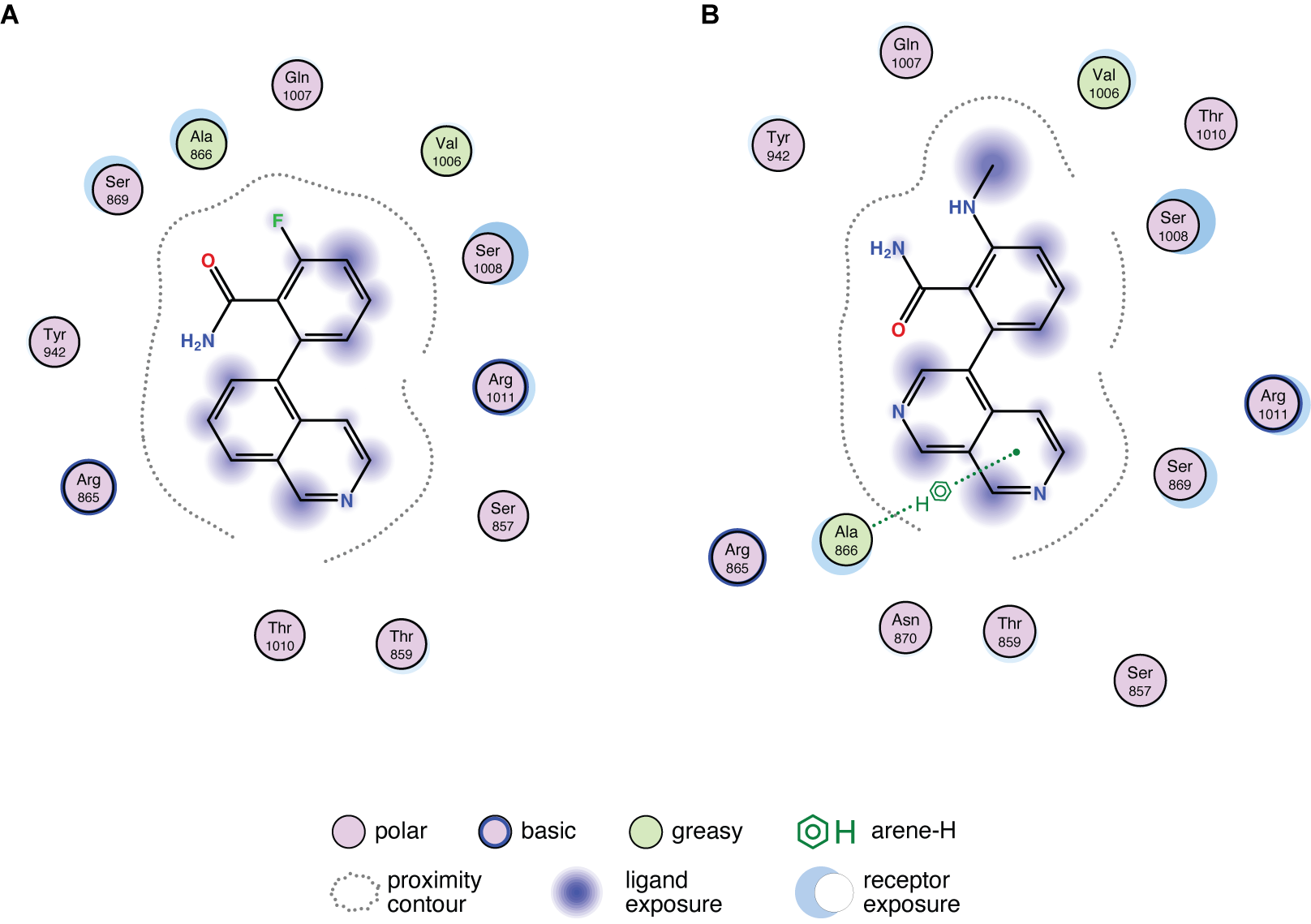

**Figure S13. 2D docking poses.** **A,B)** 2D-diagram of predicted top-scoring ligand interaction generated with MOE for GHP-88309 (A) and GHP-88310 (B). The predicted arene-hydrogen bond interactions between the isoquinoline ring of GHP-88310 and L residue A866 is shown as dotted green line.

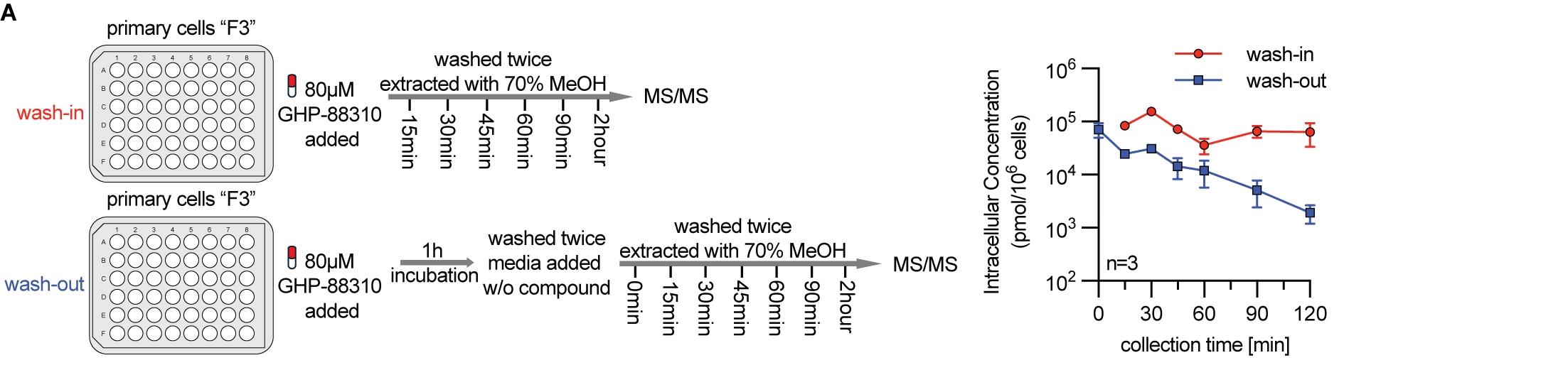

**Figure S14. Cellular uptake and retention of GHP-88310. A)** Cellular uptake and retention in undifferentiated HBTECs (female donor, F3). Symbols represent means ± SD.

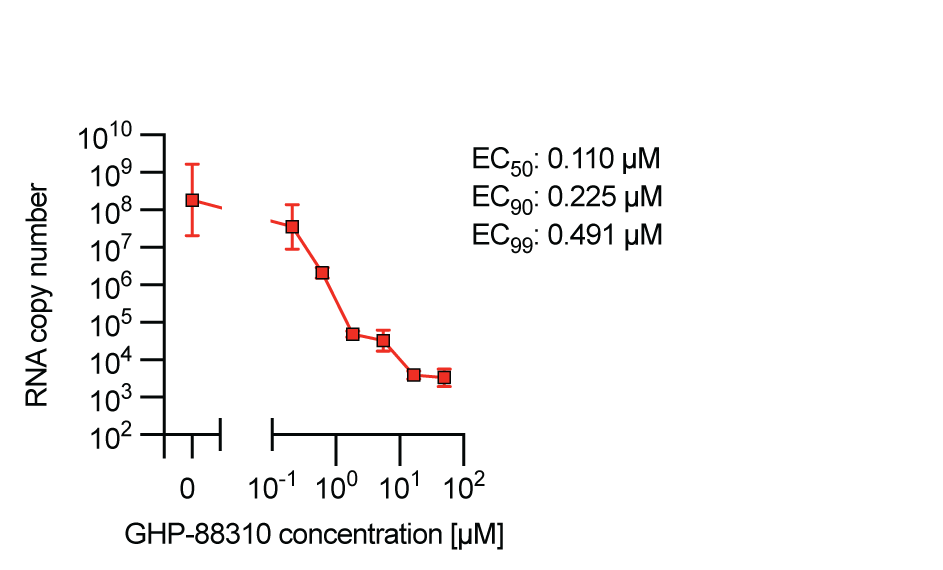

**Figure S15. Quantitation of HPIV3 RNA copy numbers through qRT-PCR.** Viral RNA copy numbers detected in samples of the dose-response assay shown in Fig. 6D. Active concentrations were calculated based on 4-parameter variable slope non-linear regression models. Symbols indicate geometric means ± geometric SD, lines connect geometric means; n=3.

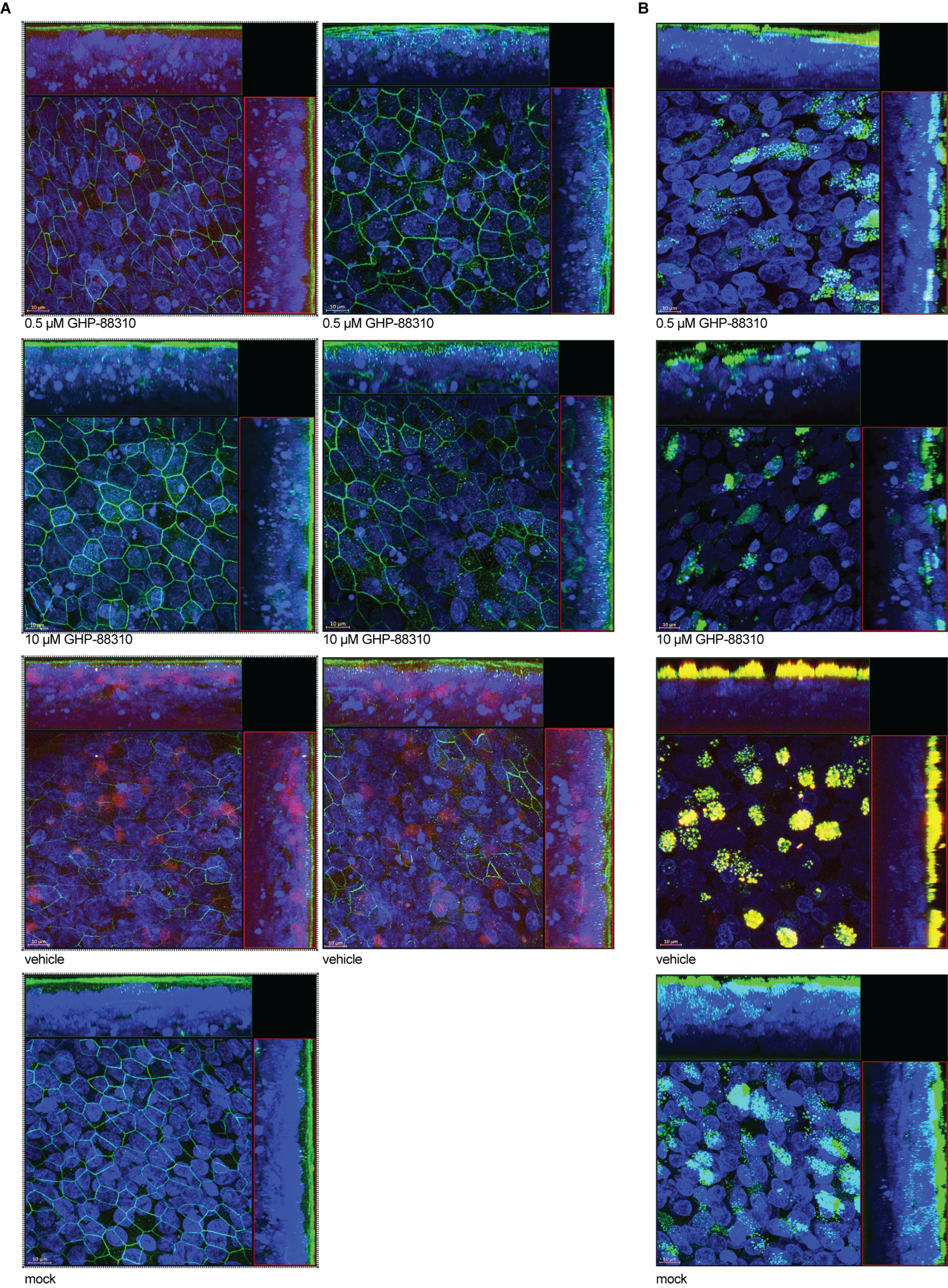

**Figure S16. Confocal imaging of GHP-88310 antiviral activity in human airway epithelium organoids. A,B)** Confocal microphotographs of organoids infected with HPIV3 were taken 2 dpi. Z-stacks of 100-150 0.22 µm slices with 63× oil objective. Scale bar, 10 µm. In (A), cells were stained with anti-PIV3 HN (red), anti-ZO-1 (tight junctions; green), and Hoechst 34580 (nuclei; blue). Images shown in (Figure 6G) are highlighted by a dashed box. In (B), cells were stained with anti-PIV3 HN (red), anti-MUC5AC (mucin; green), and Hoechst 34580 (nuclei; blue).

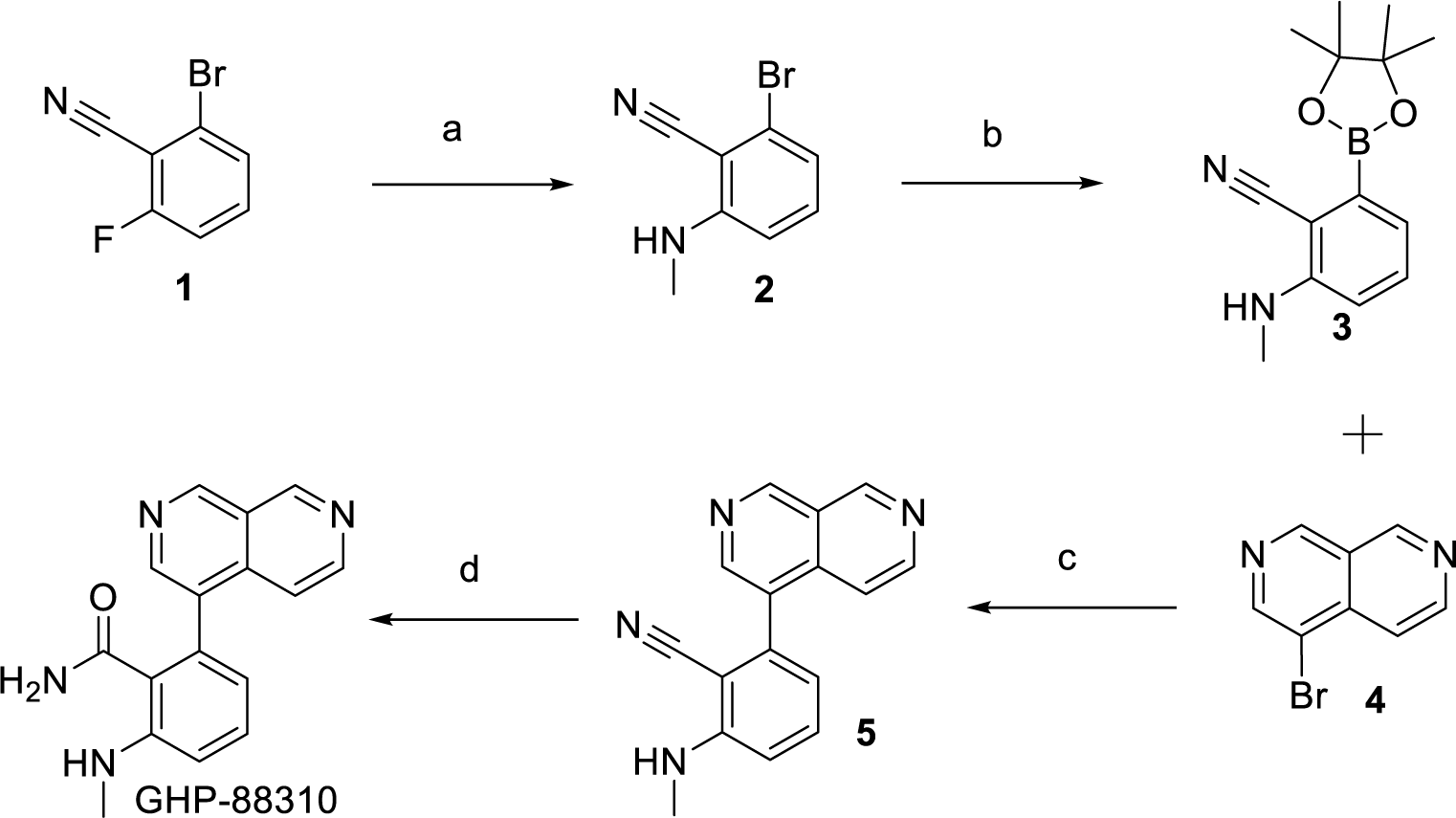

a, MeNH_2_, MeCN, 60°C, 24 hrs

b, B_2_Pin_2_, PD(dppf)Cl_2_ CH_2_Cl_2_, KOAc, 1,4-dioxane, 100°C, 16 hrs

c, XPhos Pd G3, K_3_PO_4_, 2-Me-THF, H_2_O, 60°C, 22 hrs

d, H_2_O_2_, DMSO, H_2_O, NaOH

**Figure S17. Synthesis approach for GHP-88310 and its analogs.** All materials were obtained from commercial suppliers and used without purification, unless otherwise noted. Dry organic solvents, packaged under nitrogen in septum-sealed bottles, were purchased from EMD Millipore and Sigma-Aldrich Co. Reactions were monitored using EMD silica gel 60 F_254_ TLC plates or an Agilent 1260 Infinity II series LC system with a diode array detector and an Agilent InfinityLab LC-MS detector. Compound purification was accomplished by liquid chromatography on a Teledyne CombiFlash NEXTGEN 300+ flash chromatography system. Nuclear magnetic resonance (NMR) spectra were recorded on a Bruker Ascend NMR spectrometer (400 MHz) at room temperature. Chemical shifts are reported in ppm relative to residual solvent signal.

For synthesis, 2-bromo-6-fluorobenzonitrile (Compound **1**, 15.0 g, 75 mmol, 1.0 eq) was charged to a 250 ml RBF followed by 150 ml of MeCN. With stirring, 33 wt% methylamine in absolute ethanol (24 mL, 3.21 eq,) was added to mixture and stirred at 60 ± 2°C for 24 hours. After completion, the mixture was cooled to 5°C. The resulting solid was collected by vacuum filtration. The filtrate was concentrated under reduced pressure to a solid mass. All solid materials collected were pooled and charged to RBF with 500 ml water and stirred for 2 hours. The colorless solid was filtered under vacuum and washed with water. The solid was dried in vacuum oven at 40 ± 5°C to get 14.5 gm (91% yield) of compound **2**.

Compound **2** (10.00 gm, 47 mmol, 1.0 eq,) was charged to a 250 ml RBF. B_2_Pin_2_ (14.5 gm, 57 mmol, 1.20 eq), KOAc (14.0 gm, 142 mmol, 3.00 eq) were added to the flask followed by the addition of 1,4-dioxane (200 ml) and Pd(dppf)Cl_2_ (2.0 gm, 2.3 mmol, 0.05 eq). The reaction mixture was degassed with Nitrogen for 15 minutes. The reaction mixture was stirred at 100 ± 5°C for 16 hours under nitrogen atmosphere. After completion, the reaction mixture was cooled and filtered through a Celite bed (50.0 gm). The solid was washed with 100 ml of 1,4-dioxane and collected filtrates were pooled and concentrated under reduced pressure. The residue was transferred to a 500 ml RBF using 2:8 EtOAc/*n*-Heptane (200 ml). The solution was treated with activated carbon and SiliaMetS DMT. The mixture was stirred at 40-45°C for 1 hour, cooled to room temperature and filtered through a Celite bed (50 gm). The solid cake was washed with 2:8 EtOAc/*n*-Heptane (100 ml). The filtrate was concentrated under reduced pressure, and the crude solid product was taken in a 250 ml RBF and treated with *n*-heptane (100 ml). The mixture was stirrer at 75 to 80°C for 1 hour. The mixture was slowly cooled to 25°C and continued stirring overnight. Next day the slurry was cooled to 5°C and filtered under vacuum. The resulting colorless solid was washed with *n*-heptane and dried in vacuum oven to get 9.5 gm (74% yield) of compound **3**.

4-bromo-2,7-naphthyridine (Compound **4**, 5.00 gm, 24 mmol, 1.0 eq), and compound **3** (7.0 gm, 1.1 eq) were charged to a 250 ml three neck RBF. 2-Me-THF (100 ml) was added to the mixture and stirred. Water (25 ml) and K_3_PO_4_ (10 gm, 2.0 eq) were added in sequence, and the mixture was purged with nitrogen thrice. After charging with XPhos Pd G3 (1.0 gm, 0.05 eq) the mixture was nitrogen purged three more times. The reaction mixture was stirred at 60 ± 2°C for 22 hours. After completion, the reaction mixture was cooled to 25°C and filtered under vacuum. The solid was washed with (5 mL) water. The organic layer from the filtrate was separated and dried. Combined solid from filtration and evaporation was treated with CPME (25 ml) and stirred at room temperature for 2 hours. The solid was collected by filtration and dried to get 4.85 gm of crude intermediate **5**. It was treated with SiliaMetS DMT (0.48 gm) in a mixture of DCM (170 ml) and MeOH (20 ml). The mixture was stirred at 40°C for an hour and at room temperature overnight. The mixture was filtered through Celite and the solid was washed with 9:1 DCM/MeOH. The filtrate was evaporated to dryness and the SiliaMetS DMT treatment was repeated until desired Pd levels were achieved.

Compound **5** (2.0 g, 7.7 mmol, 1.0 eq,) was charged to a 100 ml RBF. DMSO (20 ml) and water (20 ml) were added to mixture followed by the addition of 1 N NaOH (7.7 mL, 1.00 eq). The mixture was stirred well and cooled. After cooling to 5°C, 50% H_2_O_2_ (1.0 g, 2.00 eq) was added slowly over a period of 5 minutes, maintaining the temperature below 10°C. The reaction mixture was stirred at room temperature. Next day another lot of 50% H_2_O_2_ (0.30 g) was added. After stirring for another 16 hours, water (60 ml) was added to the mixture and stirred. Solid was collected by filtration and washed with water. The crude solid was charged to a 100 ml RBF after drying to which DMF (5.5 ml) was added and heated 70-75°C for 2 hours. The mixture was slowly cooled and maintained at 0-5°C, for 2 hours. The solid was filtered and washed with cold methanol (3 ml). The colorless solid was collected and dried in oven at 45-50°C to get EIDD-3608 as colorless solid 1.63 gm (75% yield).

Other analogs GHP-88369, GHP-88370, GHP-88382, GHP-88345 and GHP-88346 were synthesized using analogous procedures and were also fully characterized (supplementary Data File S1).

**Table S1. Elemental analysis of GHP-88309 analogs.**

| **Compound ID** | **NMR Spectroscopy** |
| --- | --- |
| GHP-88310  (EIDD-3608) | ^1^H NMR (400 MHz, DMSO-d6) δ 9.56 (d, *J* = 1.0 Hz, 1H), 9.51 (d, *J* = 0.9 Hz, 1H), 8.77 – 8.48 (m, 2H), 7.53 (d, *J* = 5.9 Hz, 1H), 7.40 – 7.29 (m, 2H), 7.11 (s, 1H), 6.76 (d, *J* = 7.4 Hz, 1H), 6.57 (d, *J* = 7.5 Hz, 1H), 5.47 (q, *J* = 4.9 Hz, 1H), 2.81 (d, *J* = 5.0 Hz, 3H). ^13^C NMR (101 MHz, DMSO) δ 170.0, 153.3, 152.5, 147.2, 147.1, 146.6, 137.1, 133.1, 131.6, 130.2, 124.0, 123.3, 118.6, 118.3, 110.4, 30.5. |
| GHP-88369  (EIDD-3569) | ^1^H NMR (400 MHz, DMSO) δ 9.4 (s, 1H), 9.0 (d, *J* = 4.4 Hz, 1H), 8.5 (s, 1H), 7.9 (s, 1H), 7.6 (d, *J* = 4.4 Hz, 1H), 7.5 (d, *J* = 5.7 Hz, 1H), 7.3 (d, *J* = 8.4 Hz, 1H), 7.3 (d, *J* = 2.6 Hz, 1H), 7.3 (s, 1H), 7.2 (dd, *J* = 8.4, 2.7 Hz, 1H), 3.9 (s, 3H); ^13^C NMR (101 MHz, DMSO) δ 169.6, 159.7, 153.9, 152.2, 147.4, 143.9, 143.2, 138.9, 132.6, 130.7, 127.0, 125.5, 118.7, 115.8, 114.0, 56.0; MS (ES-API) [M+1]^+^: 280.2. |
| GHP-88370  (EIDD-3570) | ^1^H NMR (400 MHz, DMSO) δ 9.6 (s, 1H), 9.5 (s, 1H), 8.7 (d, *J* = 5.9 Hz, 1H), 8.6 (s, 1H), 7.8 (s, 1H), 7.4 (d, J = 5.9 Hz, 1H), 7.3 (d, *J* = 8.4 Hz, 1H), 7.2 (d, *J* = 2.6 Hz, 1H), 7.2 – 7.2 (m, 2H), 3.9 (s, 3H); ^13^C NMR (101 MHz, DMSO) δ 169.0, 159. 5, 153.4, 152.2, 147.2, 146.3, 139.6, 137.4, 133.2, 131.5, 125.8, 123.3, 117.8, 115.7, 113.9, 56.0; MS (ES-API) [M+1]^+^: 280.2. |
| GHP-88382  (EIDD-3482) | ^1^H NMR (400 MHz, DMSO) δ 9.6 (d, *J* = 1.1 Hz, 1H), 9.5 (d, *J* = 1.0 Hz, 1H), 8.7 (d, *J* = 5.9 Hz, 1H), 8.7 (s, 1H), 7.7 (d, *J* = 1.6 Hz, 1H), 7.5 – 7.4 (m, 3H), 7.2 (d, *J* = 2.4 Hz, 1H), 7.2 (ddd, *J* = 7.0, 1.8, 0.6 Hz, 1H), 2.4 (s, 3H); ^13^C NMR (101 MHz, DMSO) δ 170.3, 153.3, 152.8, 147.3, 146.8, 139.9, 137.3, 134.6, 132.2, 130.7, 130.5, 128.6, 128.4, 123.3, 118.3, 19.7; MS (ES-API) [M+1]^+^: 264.1. |
| GHP-88345  (EIDD-3545) | ^1^H NMR (400 MHz, DMSO) δ 9.5 (s, 1H), 9.1 (d, *J* = 4.4 Hz, 1H), 8.6 (s, 1H), 7.7 (s, 1H), 7.6 (d, *J* = 4.5 Hz, 2H), 7.5 (s, 1H), 7.2 (d, *J* = 7.9 Hz, 1H), 6.9 (d, *J* = 8.0 Hz, 1H), 6.2 (s, 2H); ^13^C NMR (101 MHz, DMSO) δ 165.4, 153.8, 152.2, 148.4, 146.8, 145.6, 143.8, 130.6, 128.2,125.6, 124.7, 120.0, 109.4, 102.5; MS (ES-API) [M+1]^+^: 294.1. |
| GHP-88346  (EIDD-3546) | ^1^H NMR (400 MHz, DMSO) δ 9.6 (d, *J* = 1.0 Hz, 1H), 9.5 (d, *J* = 1.0 Hz, 1H), 8.7 (d, *J* = 6.0 Hz, 1H), 8.6 (s, 1H), 7.7 (s, 1H), 7.5 (d, *J* = 5.9 Hz, 1H), 7.4 (s, 1H), 7.1 (d, *J* = 7.9 Hz, 1H), 6.9 (d, *J* = 7.9 Hz, 1H), 6.2 (d, *J* = 4.0 Hz, 2H); ^13^C NMR (101 MHz, DMSO) δ 165.7, 153.4, 152.4, 148.1, 147.3, 146.4, 145.4, 137.5, 130.9, 127.1, 125.1, 123.3, 120.8, 117.9, 109.3, 102.4; MS (ES-API) [M+1]^+^: 294.1. |

**Table S2. GHP-88310-class resistance sites.** All sequences were aligned to MeV. GHP-88310-class resistance sites in are highlighted in purple (MeV), blue (HPIV3), and green (SeV), respectively. Orange highlights depict resistance sites that emerged in both MeV and SeV.

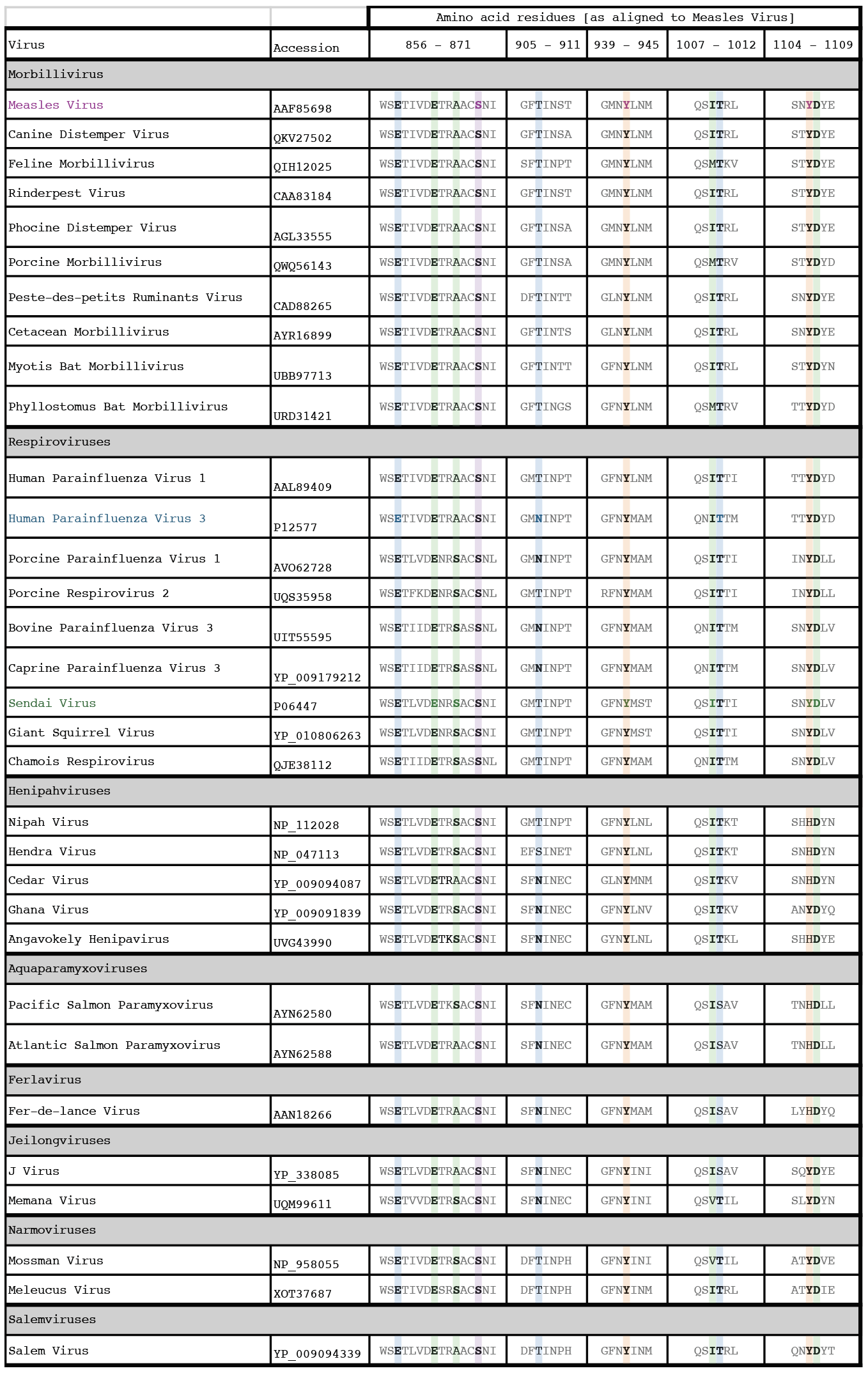
